# One species or several? Genomic evidence for deep divergence within the widely distributed anemonefish *Amphiprion clarkii* species complex

**DOI:** 10.64898/2026.09.17.752292

**Authors:** Jann Zwahlen, Anna Marcionetti, Manon Mercader, Saori Miura, Carlos Leiva, James Davis Reimer, Nicolas Salamin, Vincent Laudet

## Abstract

Defining what a species is remains one of the biggest questions in biology and directly challenges species delimitation in species complexes. The wide-spread Clark’s anemonefish *Amphiprion clarkii* has the widest range of all anemonefishes, from the Indian Ocean to the Central Pacific. However, the taxonomy of this species has been confused since its initial description due to its great variation in morphology and pigmentation even within the same locality. Here, we combined whole genome re-sequencing data from 166 individuals spanning most of the distribution of *A. clarkii,* including remote islands such as Ogasawara and Guam in the northwestern Pacific, and also including three individuals of the closely related *A. tricinctus* from the Marshall Islands. Our population genomic analyses reveal an unexpected level of divergence, with five deeply separated lineages corresponding to the Indian Ocean, Central Indo-Pacific, Melanesia, and the isolated population of Ogasawara. The fifth lineage comprises *A. clarkii* from Guam and *A. tricinctus*, revealing that *A. tricinctus* is phylogenetically nested within *A. clarkii*, challenging its current taxonomic status. The most divergent lineage, Melanesia, is genetically more distant from other *A. clarkii* lineages than several recognised species pairs within the *Amphiprion* genus, yet demographic modelling shows ongoing gene flow between all lineages. Together, our results highlight a hidden diversity and stress the necessity of an integrative taxonomic revision of the *A. clarkii* species complex.

## Introduction

Few concepts in biology have been as debated, and as fundamental, as that of the species itself (Coyne & Orr, 2004). This has direct implications for species delimitation (De Queiroz, 2007), the process of evaluating boundaries between populations or species. Originally, species description heavily relied on morphological features of individuals, however, supplementary molecular phylogenetic approaches have become recommended (Dayrat, 2005; Jörger & Schrödl, 2013; Padial et al., 2010). The emergence of genomic data has further improved the power of molecular data to identify cryptic divergence not readily visible from morphological features, such as meiofaunal slugs from the genus *Pontohedyle* (Jörger & Schrödl, 2013) or the blenniid *Cirripectesalboapicalis* species complex (Delrieu-Trottin et al., 2018). The shift in focus from morphological to genomic and molecular data has raised concerns of over-splitting species (Coates et al., 2018; Isaac et al., 2004), especially when relying only on one method (Carstens et al., 2013; Chambers & Hillis, 2020). However, especially in the case of cryptic species which are characterised by not having clear morphological differences, multiple approaches for an integrative taxonomy may not always be possible (Bickford et al., 2007; Jörger & Schrödl, 2013).

This challenge is particularly pronounced in marine ecosystems, which have long been considered highly connected because of the apparent absence of physical barriers to dispersal, a view further supported by the large population sizes and extensive connectivity observed in many marine species (Carrete Vega & Wiens, 2012; Palumbi, 1994). Indeed, Briggs’ (1960) list of 107 circumtropical fish species was later expanded to nearly 300 species (Gaither et al., 2016). However, almost three-quarters of these species have not been studied adequately using genetic markers, and more than 10% exhibit unclear delimitation, revealing substantial cryptic diversity within circumtropical fishes (Gaither et al., 2016; Nyegaard et al., 2018; Sawai & Nyegaard, 2022; Underkoffler et al., 2018). Despite high dispersal potential, the ocean is not an unstructured environment; historical events, temperature and salinity gradients and fronts, and oceanic currents (Belkin et al., 2009) act as barriers to gene flow, shaping fine-scale population structure. In many marine taxa, a biphasic life cycle with a pelagic larval phase facilitates dispersal across vast distances; however, pelagic larval duration, behaviour, and oceanographic context can strongly modulate connectivity, sometimes restricting gene flow even across short spatial scales (Cowen & Sponaugle, 2009). These patterns highlight that, even in a seemingly continuous marine realm, biodiversity and speciation are governed by a complex interplay between dispersal and environmental heterogeneity.

Even within a species, such factors can influence genetic diversity, which is thought to follow two related patterns. According to the central-marginal hypothesis (Antonovics, 1976; Brussard, 1984), central populations have higher genetic diversity than peripheral ones. Similarly, isolated populations, such as island populations which are often considered marginal, are expected to have lower genetic diversity than well connected populations (e.g. on mainland; Frankham, 1997). A review of over a hundred studies found that although two-thirds of them showed reduced genetic diversity towards the periphery, the differences were often comparatively small (Eckert et al., 2008), and a more recent analysis found only half of examined studies with this pattern (Pironon et al. 2017). For example, isolated angelfish populations of both widespread and endemic species retained high genetic diversity (Hobbs et al., 2013). Therefore, neither peripheral position nor isolation are necessarily linked with lower genetic diversity in marine fishes.Anemonefishes from the genus *Amphiprion* provide a great opportunity to study species and population structure mechanisms due to their short larval duration of only around two weeks, which limits their ability to disperse (Jones et al., 2022). The genus currently consists of 29 species distributed in the tropical to subtropical Indo-Pacific, all living in obligate symbiosis with giant sea anemones (Fautin & Allen, 1992; Laudet & Ravasi, 2022; O’Donnell et al., 2025). *Amphiprion* species differ in the number of associated host anemones and distribution (Litsios et al., 2014a). Recent studies suggest their diversification is driven by more than one factor: host identity shapes shared colour patterns among species using the same anemones (Gaboriau et al., 2025), while separate ecological strategies, such as differences in swimming, metabolism, and host dependence, also drive diversification, independent of host use (Mercader et al., 2025). In addition, growing evidence suggests that the current taxonomy of anemonefishes underestimates their true diversity, with several nominal species likely representing complexes of cryptic or incipient lineages. For example, *A. chrysopterus* was recently split into *A. chrysopterus* and *A. maohiensis* (O’Donnell et al., 2025), and *A. polymnus* has been proposed to be a complex of up to three different species (Fitzgerald et al., 2026). Similar patterns have been revealed in *A. ocellaris*, where population genomic studies have identified strong genetic structuring and restricted gene flow across the Indo-Malay Archipelago (Timm & Kochzius, 2008), and in the skunk anemonefishes, which have been shown to contain several deeply divergent lineages shaped by historical barriers and local adaptation (Marcionetti et al., 2024).

The Clark’s anemonefish, *Amphiprion clarkii*, illustrates both the taxonomic ambiguity and striking phenotypic plasticity that can characterise some species of *Amphiprion*. Geographically, *A. clarkii* spans the widest range of any anemonefish, latitudinally from central Japan to Australia and longitudinally from Oman to New Caledonia, and is uniquely associated with all ten currently valid host anemone species (Fautin & Allen, 1992). Historically, this complex grouped *A. clarkii* with relatively morphologically similar species such as *A. tricinctus*, *A. chrysopterus*, and *A. chagosensis* (Allen, 1972), but molecular phylogenies have demonstrated that these species were not all closely related (Laudet & Ravasi, 2022; Quenouille et al., 2004; Santini & Polacco, 2006). Nevertheless, the term “*clarkii* complex” has been used to describe the exceptional diversity within *A. clarkii* itself (e.g. Rowlett, 2020), which may in fact encompass several cryptic lineages. Indeed, mitochondrial analyses have placed *A. tricinctus,* a species not overlapping geographically with *A. clarkii,* within the *A. clarkii* clade (Litsios et al., 2014b; Litsios & Salamin, 2014), and the maximum genetic distance within *A. clarkii* exceeds that between *A. clarkii* and *A. tricinctus* (He et al., 2022), raising questions about the currently accepted species boundaries. Across its wide range, populations exhibit remarkable morphological variation, especially at the pigmentation level. This is further complicated by phenotypic plasticity linked to sea anemone host association, which occurs across most of the species’ range. Individuals associated with *Stichodactyla* are typically black, while those in *Radianthus, Heteractis* and *Entacmaea* are more orange (Fig. 1A; Militz et al., 2016), and additional local traits such as body shape, size, and pigmentation, further diversify the phenotype (Bell et al., 1982). In the isolated Ogasawara Islands, for instance, *A. clarkii* are entirely black except for their white bars (Bell et al., 1982; Moyer, 1976), despite associating only with *E. quadricolor*. Allen (1972) also noted pigmentation differences between Indian and Pacific Ocean populations. However, without accounting for host identity, interpretations remain uncertain. Geographic variation extends to tail colouration and meristic traits, such as spine numbers, across Japanese populations (Bell et al., 1982; Moyer, 1976). Recently, Schmid et al. (2024) showed that *A. clarkii* consists of four lineages: a well-connected central lineage containing populations from Indonesia, the Philippines and Taiwan, a Pacific Ocean lineage containing populations from Papua New Guinea (PNG) and the Solomon Islands, a lineage in New Caledonia, and a fourth lineage comprising the Maldivian population. Despite their strong separation, these populations still have ongoing gene flow. Furthermore, the authors hypothesised a stepping-stone model for dispersal, with an origin in the central Indo-Pacific (Schmid et al., 2024). Similarly, Clark et al. (2021) also showed three main clades (without New Caledonia). Together, these observations highlight *A. clarkii* as a mosaic of divergent populations with high phenotypic plasticity and unresolved taxonomy.

**Figure 1.**
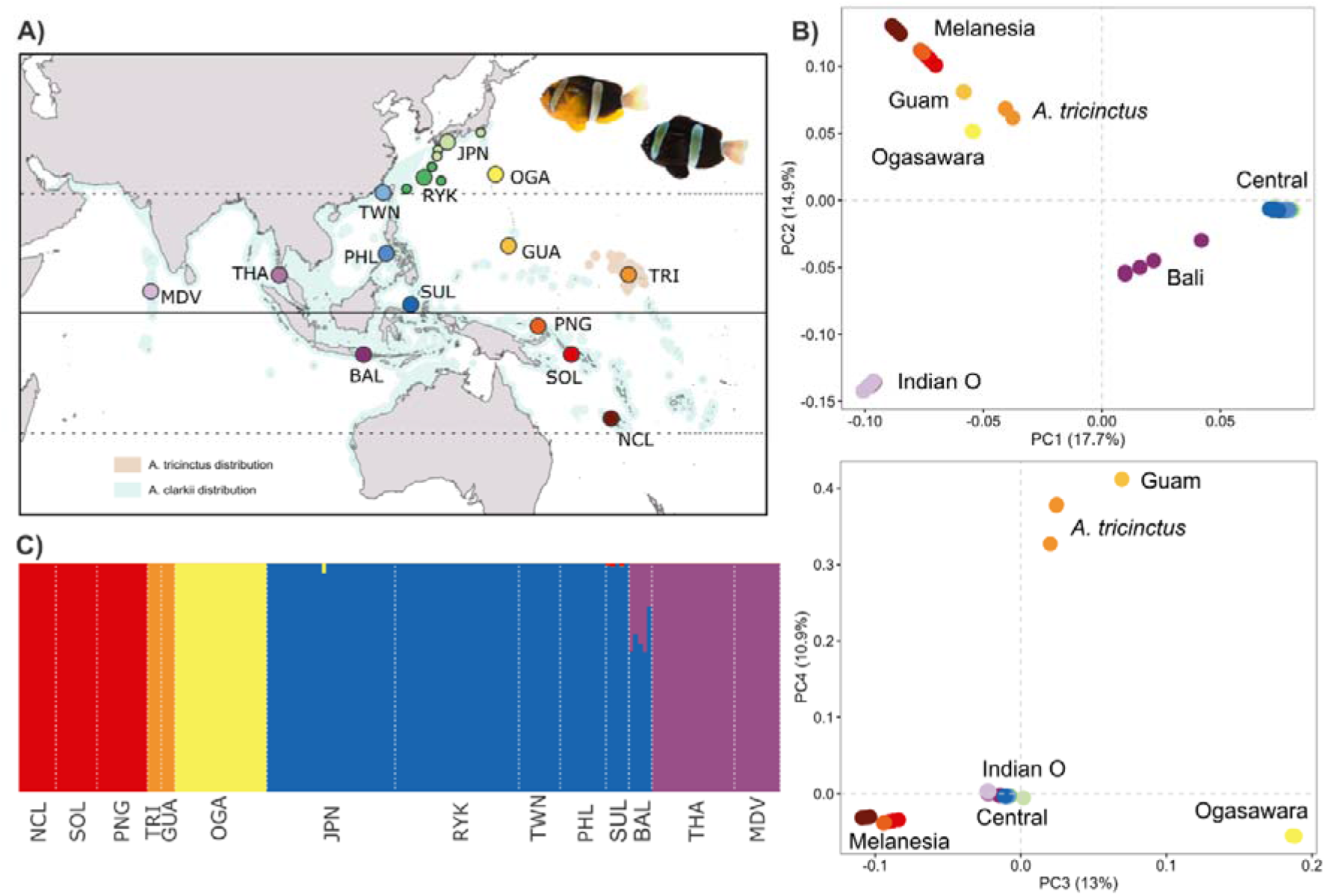
Sampling locations and genetic structure of *A. clarkii* populations. **A)** Map showing the locations of the samples. Shaded blue and orange colour is the distribution of *A. clarkii* and *A. tricinctus* from IUCN Red List (Allen et al., 2022; Jenkins et al., 2017). MDV Maldives; THA Thailand; BAL Bali (Indonesia); SUL Sulawesi (Indonesia); PHL Philippines; TWN Taiwan; RYK Ryukyus (smaller dots represent our original sampling locations); JPN mainland Japan; OGA Ogasawara; GUA Guam, TRI *A. tricinctus*; PNG Papua New Guinea; SOL Solomon Islands; NCL New Caledonia. The black and orange colour morphs from Okinawan *A. clarkii* are shown in the top right corner. **B)** First four principal components of PCA from the SNP dataset showing the diversity of *A. clarkii* populations and revealing that *A. tricinctus* lies within *A. clarkii*, most closely to the Guam population. Number in parentheses is % of variance explained. Dashed lines represent 0 coordinates. **C)** Result from admixture analysis based on our SNP dataset with *K=5* highlighting the five main clusters in the *A. clarkii* species complex.

Here, we build on Schmid et al. (2024) and the previously described *A. clarkii* population structure with strong divergence but ongoing gene flow. We have added samples from the northern range limit (Japan) and very isolated islands (Ogasawara and Guam), as marginal and isolated groups are often overlooked in population studies. With our expanded dataset we investigated whether small, isolated populations follow similar trajectories to those of major diverged regions, and if these northern populations are an extension of the well-mixed central Indo-Pacific region. To bring the high divergence within *A. clarkii* (He et al., 2022; Schmid et al., 2024) into a wider perspective for potential species delimitation, we also included *A. tricinctus* samples as the most closely related species (Litsios & Salamin, 2014) in our analyses.

## Materials & Methods

### Sample collection, DNA extraction and sequencing

We caught wild adult *A. clarkii* around Japan and Guam (all required permits obtained) using hand nets and collected small fin clips. Once on land, the fin clips were transferred into tubes with RNAlater and stored at 4°C for several days and then at -20°C until DNA extraction. Furthermore, we collected fin clips from captive *A. tricinctus* originally from Marshall Islands and now kept at Academia Sinica’s Marine Research Station in Yilan, Taiwan, and received tissue samples from additional Japanese populations from the National Museum of Nature and Science of Japan. DNA was extracted using Promega Maxwell Blood DNA Extraction Kit (Catalog# AS1400). Paired-end libraries (Library Prep Kit for Illumina (Catalog# E7430)) were prepared for each sample, and the samples sequenced using DNA shotgun sequencing on NovaSeq 6000 or NovaSeq X platforms with 300 cycles and a read-length of 150 basepairs. We aimed for 136 million reads per sample, expecting an average genome-wide depth of 15X. Furthermore, we downloaded samples from worldwide *A. clarkii* populations (García Jiménez et al., 2025, Schmid et al., 2024) from NCBI, resulting in the most complete dataset yet assembled for this species complex with 166 samples spanning across 20 populations including *A. tricinctus* (Table S1.; Fig. 1A). Additionally, we added one sample of *A. chrysopterus* as an outgroup for later analyses.

### Processing of sequencing data and genotype likelihood calculation

Once sequencing was completed, we removed the adapters, low-quality basepairs at the ends of reads, and reads shorter than 36 basepairs using trimmomatic/0.39 (Bolger et al., 2014). We assessed read quality before and after trimming using fastqc (Andrews, 2010). Reads were aligned to the *A. clarkii* reference genome (Moore et al., 2023) using the bwa-mem algorithm of the Burrows-Wheeler Aligner (Li, 2013). Samtools was used to transform sam to bam files and to index the final files (Danecek et al., 2021), picard was used to sort the bam files, mark and remove duplicates and to replace the read groups (https://broadinstitute.github.io/picard/). Since we obtained our sequences from multiple sequencing runs, we calibrated the sequences to reduce sequencing bias. For this, we created a vcf file with high-confidence SNPs from a single sequencing run with 60 samples from Japan: First, we ran the steps as described above, and continued to call haplotypes, combine vcfs, genotype the variants and filter the genotyped reads (QD < 2, SOR > 3, FS > 60, MQ < 40, MQRankSum < -12.5, ReadPosRankSum < -8.0) using GATK (Van der Auwera & O’Connor, 2020). We then used GATKs BaseRecalibrator and ApplyBQSR for a first calibration and repeated the whole steps on the same samples to get a filtered vcf file that was calibrated twice. We then used this file as baseline for the calibration of the bam files of all samples. Due to uneven sequencing depths and population sizes, genotype likelihoods were estimated using ANGSD (Korneliussen et al., 2014). We calculated the genotype likelihood on all sites using quality filters -minMapQ 30, -minQ 20, -baq 1 and called posterior probabilities (-doPost 1) and called genotypes (-doGeno 32) to get a bcf file with hard-called genotypes. Since the dataset was too large to process at once, we did this for each chromosome separately, and used bcftools (Danecek et al., 2021) to combine the chromosomes, convert the files to vcf, and apply shallow filtering, retaining sites with MAC ≥ 2 and ≤ 10% missing genotypes.

### Population genetics and structure

To understand the general population structure in our study, we used an LD-pruned dataset (PLINK --indep-pairwise 50 5 0.2 (Purcell et al., 2007); 13,319,227 SNPs) to infer population structure in ADMIXTURE 1.3 (Alexander et al., 2009). We ran ADMIXTURE from *K*=*1* to *K*=*12* and picked the best number of clusters based on the lowest cross validation error (Fig. S1). Furthermore, we did a principal component analysis (PCA) using PLINK (Purcell et al., 2007), and visualised the resulting principal components using ggplot2 (Wickham, 2011) and plotly (Sievert, 2020) in R (R Core Team, 2022).

Furthermore, we used common population genetics indices such as fixation index (*F*_ST_) to measure relative differentiation between populations, the absolute divergence (*d*_xy_) and *π*, to measure the nucleotide diversity for each population. We calculated these indices (*F*_ST_, *d*_xy_ and *π*) using sliding windows of 5,000 SNPs with the script popgenWindows.py from https://github.com/simonhmartin/genomics_general/.

As we originally focused on the Japanese populations, our sampling across Japan was geographically more widespread and extensive than the sequences retrieved from NCBI. Based on an initial admixture analysis (Fig. S2) and geography, we grouped the Japanese samples into two ensembles: mainland Japan (Kochi, Yakushima, Kagoshima, Shikine) and the Ryukyus (Ishigaki, Okinawa, Aka, Amami and Daito) for consistency with the other locations.

To test for relatedness among individuals, we calculated the pairwise kinship coefficient from our unpruned dataset using the kinship option in KING (Manichaikul et al., 2010). The pairwise kinship inference using the kinship option is robust to population structure, and the coefficient values indicate the degree of relatedness (Manichaikul et al., 2010). Furthermore, we also searched for Runs of Homozygosity (RoH) as an indicator for past bottlenecks or recent inbreeding. We used PLINK (Purcell et al., 2007) on our unpruned dataset and searched in sliding windows of 300kb. To call a RoH, we required at least 50 SNPs and a minimum density of 1 SNP per 50kb, but allowed up to three heterozygous sites per sliding window for potential genotyping errors, and required at least a 1000kb gap between two windows, as described in Foote et al. (2021).

### Phylogenetic trees

We constructed a phylogenetic tree from a reduced SNP dataset to reduce the computational demands. First, we selected 5% random SNPs from our dataset using vcflib’s vcfrandomsample (Garrison et al., 2022) and transformed the reduced vcf to fasta using vcf2phylip (Ortiz, 2019). We used IQtree2 (Minh et al., 2020) to remove invariant sites (e.g. ambiguously constant sites for which the alternative alleles are only present in a heterozygous state), and then used IQtree2’s ModelFinder (Kalyaanamoorthy et al., 2017) to select our model to infer the tree. The best model was TVM+F+ASC+R5, a transversion model with equal transition rates but unequal base frequencies (TVM), empirical base frequencies (F), ascertainment bias correction (ASC) to account for only variable sites (Lewis, 2001) and five categories of substitution rates with the FreeRate model R5 (Soubrier et al., 2012; Yang, 1995). We calculated SH-like approximate likelihood ratio test (SH-aLRT) using 1000 replications (Guindon et al., 2010) and did 1000 ultra-fast bootstraps for the branch support directly (Hoang et al., 2018). We used figtree v.1.4.4 (https://tree.bio.ed.ac.uk/software/figtree) to visualise the tree and re-rooted it using *A. chrysopterus* as outgroup.

In addition to the full genome, we produced trees using all SNPs for each chromosome separately, with the standard settings for SNP data, GTR+ASC. These chromosome-scale trees allowed us for general check of phylogenetic discordance within the full genome. We reduced the chromosomal trees to one individual per main clade to obtain the main topologies and compared them with each other and the whole-genome tree by calculating pairwise Robinson-Foulds distances using the ape package in R (Paradis & Schliep, 2019). We generated plots in ggplot2 (Wickham, 2011), using classical multidimensional scaling (cmdscale) for the topological distances.

### Demographic history

The demographic history of *A. clarkii* follows the stepping stone model, where the Central population is the species’ origin, with Indian Ocean and Melanesian populations emerging from there (Schmid et al., 2024). To investigate the demographic history of the remote populations of Ogasawara and Guam, as well as the positioning of *A. tricinctus* within *A. clarkii*, we used fastsimcoal2, a fast, sequential Markovian coalescent to simulate historic events (Excoffier et al., 2021). Fastsimcoal2 requires a site frequency spectrum (SFS) format, which we created with easySFS from our LD-pruned dataset (Gutenkunst et al., 2009). To reduce computational demands and the variation within populations, we excluded New Caledonian individuals from Melanesia and Bali individuals from the Central group. For Ogasawara we considered the following scenarios (Fig. S3A) based on the chromosomal nuclear phylogenetic trees (Fig. S4B/C): 1) Ogasawara and Melanesia splitting off independently from the Central population, and 2) Melanesia diverging from the central population, and then Ogasawara diverging from Melanesia. For both scenarios we had three migration scenarios: a) ongoing and past gene flow between all populations, b) ongoing gene flow only between Melanesia and the Central population, and c) ancient gene flow between Melanesia and the Central population, but currently no ongoing gene flow (which would contradict Schmid et al., 2024). For *A. tricinctus* we wanted to test the main scenarios regarding the association with *A. clarkii*: 1) *A. tricinctus* emerged from Melanesia within the *A. clarkii* clade, 2) from the Central clade, or 3) an ancestral group split into *A. clarkii* and *A. tricinctus*, with a more recent secondary contact with introgression between the two species. Furthermore, we added 4) where Melanesia and the Central group emerged from an ancestral population, to account for potential preference of secondary contact (Momigliano et al., 2021; Smith & Hahn, 2024). In all the scenarios with ancestral populations we gave options with different migration and isolation patterns and changed the order between the emergence of Guam and *A. tricinctus* (all models in Fig. S3B).

For every scenario, we ran the model 100 times, each with 200,000 coalescent simulations and 50 ECM cycles, and ignored monomorphic sites. Since the mutation rate in *A. clarkii* is still unknown, we fixed the splitting time between central Indo-Pacific and Melanesia to 60,060 generations, based on Schmid et al. (2024). We selected the best run for each model by finding the lowest difference between the obtained likelihood and maximum possible likelihood, using a python script (Joana Meier, https://raw.githubusercontent.com/speciationgenomics/scripts/master/fsc-selectbestrun.sh). For every model’s best run we calculated the Akaike information criterium (AIC) using an Rscript from Meier et al. (2017) and the likelihood distribution using the parameters for another 100 runs to find the best scenario.

We used parametric bootstrapping in fastsimcoal2 to obtain confidence intervals for the best scenario. For this, we created 100 SFS from the maximum composite likelihood parameter estimates of the best initial run. For each of the SFS replicates we used 30 independent runs with 200,000 simulations and 20 ECM cycles (Excoffier et al., 2013; Schmid et al., 2024) to re-estimate the parameters and calculate 95% confidence intervals.

In addition to fastsimcoal2, we used Dsuite (Malinsky et al., 2021) to check for introgression between more distantly related populations not included in the demographic simulations. First, we ran Dtrios for each chromosome separately, using *A. chrysopterus* as outgroup. Next, we used the command DtriosCombine to combine the chromosomal output into one file. From this file, we calculated the f-branch statistic (*f_b_*), which highlights branches in a tree which share alleles excessively compared to what would be expected from their positions, using Dsuite’s Fbranch command. We reduced the nuclear tree into populations using the R package ape (Paradis & Schliep, 2019) by selecting the first individual per population and collapsing the remaining branches. Finally, we visualised the results using the script dtools.py, provided by Dsuite (Malinsky et al., 2021).

### Mitochondrial phylogenetic tree

We built a mitochondrial DNA (mtDNA)-based phylogenetical tree to compare it with the nuclear DNA estimates. First, we used Burrows-Wheeler Aligner to align our clean reads to the *A. clarkii* reference mitochondrial genome (Tao et al., 2016b). Next, we used bcftools for variant calling and creating a consensus sequence (Danecek et al., 2021). We used all mitochondrial protein-coding genes to create the mitochondrial tree. Additionally, we included sequences from other species pairs within *Amphiprion* based on the availability of reference mitochondrial genome, namely *A. frenatus* (Li et al., 2015), *A. ephippium* (Thongtam Na Ayudhaya et al., 2019), *A. percula* (Tao et al., 2016a), *A. ocellaris* (Mabuchi et al., 2007), *A. perideraion* (Hu et al., 2016), and *A. akallopisos* (Thongtam Na Ayudhaya et al., 2019). Additionally, we included an individual *A. clarkii* from the South China Sea (Tao et al., 2016b) as a verification of our clustering. Fasta files were aligned using the G-INS-i method in MAFFT with 1000 iterations (Katoh & Standley, 2013). We constructed the tree using the partition model -p on IQTree2 with 1000 ultra-fast bootstraps (Chernomor et al., 2016; Minh et al., 2020). We visualised the tree in figtree v.1.4.4 (https://tree.bio.ed.ac.uk/software/figtree) and rooted it from the *A. percula* and *A. ocellaris* branch, as their positioning within *Amphiprion* has been well established (Gaboriau et al., 2025; Santini & Polacco, 2006). Cophenetic distances were calculated using ape’s (Paradis & Schliep, 2019) function cophenetic.

## Results

### Sequencing and variant calling

We analysed 166 samples from a wide geographic area spanning from Japan to New Caledonia, including samples from the Indian Ocean (Fig. 1A). For the individuals sequenced in this study (Table S1), Illumina sequencing yielded an average of 158.8 million pair reads per sample (ranging from 120.7 to 213.7 million), and an average of 138.2 million (57.4 to 216.7 million) when combining our data with those from Schmid et al. (2024) and García Jiménez et al. (2025). The sequencing depth of the newly sequenced samples ranged from 13.1 – 30.7X, while some of the additional samples had a slightly lower sequencing depth (down to 5.7X).

Our analysis using the ANGSD pipeline identified 46 million single nucleotide polymorphisms (SNPs) across the dataset, which was reduced to 35.3 million SNPs after filtering.

### Population structure

We performed PCA and admixture analysis to infer the overall population structure. Looking at the PCA (Fig. 1B), major regions were separated by different principal components (PC). PC1 distinguished populations from the central Indo-Pacific (mainland Japan, Ryukyus, Taiwan, the Philippines, Sulawesi and Bali; from now referred to as Central population). PC2 separated the Indian Ocean populations (Maldives and Thailand), while the Melanesian samples (PNG, Solomon Islands and New Caledonia) were characterised by both, PC2 and PC3, the latter also clearly differentiating Ogasawara. Guam and *A. tricinctus* were separated from the rest by PC4, which explained 10.9% of the variance.

Similarly, the best *K* in ADMIXTURE was 5 (Fig. S1) and separated the four major regions (Central, Indian Ocean, Melanesia and Ogasawara) into different clusters (Fig. 1C), with Bali having some influence from the Indian Ocean. *A. clarkii* individuals from Guam and *A. tricintus* individuals were grouped within the same cluster, in line with the PCA results.

To understand the relationship between the different populations, we computed *F*_ST_, a fixation index to measure population differentiation, *d*_xy_, a measure of population divergence, and *π* for nucleotide diversity, using sliding windows along the genome. We observed that *A. clarkii* populations from the Central population had very limited structure, with *F*_ST_ values ranging from 0.005 to 0.044 (Fig. 2; above diagonal). Similarly, Solomon Islands and PNG populations presented a low *F*_ST_ (0.011). On the other hand, all other populations had relatively strong differentiation. The highest values were found for the Ogasawara population with pairwise *F*_ST_ values ranging from 0.27 to 0.53, indicating substantial structure. The population divergence (*d*_xy_) values showed a similar picture, with lower values in the Central population (Fig. 2; below diagonal), but also within Melanesia and the Indian Ocean. Overall, the highest divergence was found between the Melanesian populations and the rest, reaching up to 0.139. The nucleotide diversity *π* followed a different pattern: the Central population had the highest diversity values (0.005), while Guam, Ogasawara, PNG and Solomon Islands presented the lowest nucleotide diversity (0.002; Fig. 2; diagonal). Taken together, the central Indo-Pacific marked a region of high connectivity with low structure between populations and high nucleotide diversity, whereas the other major regions, as well as isolated populations were highlighted by strong structure and lower diversity.

**Figure 2.**
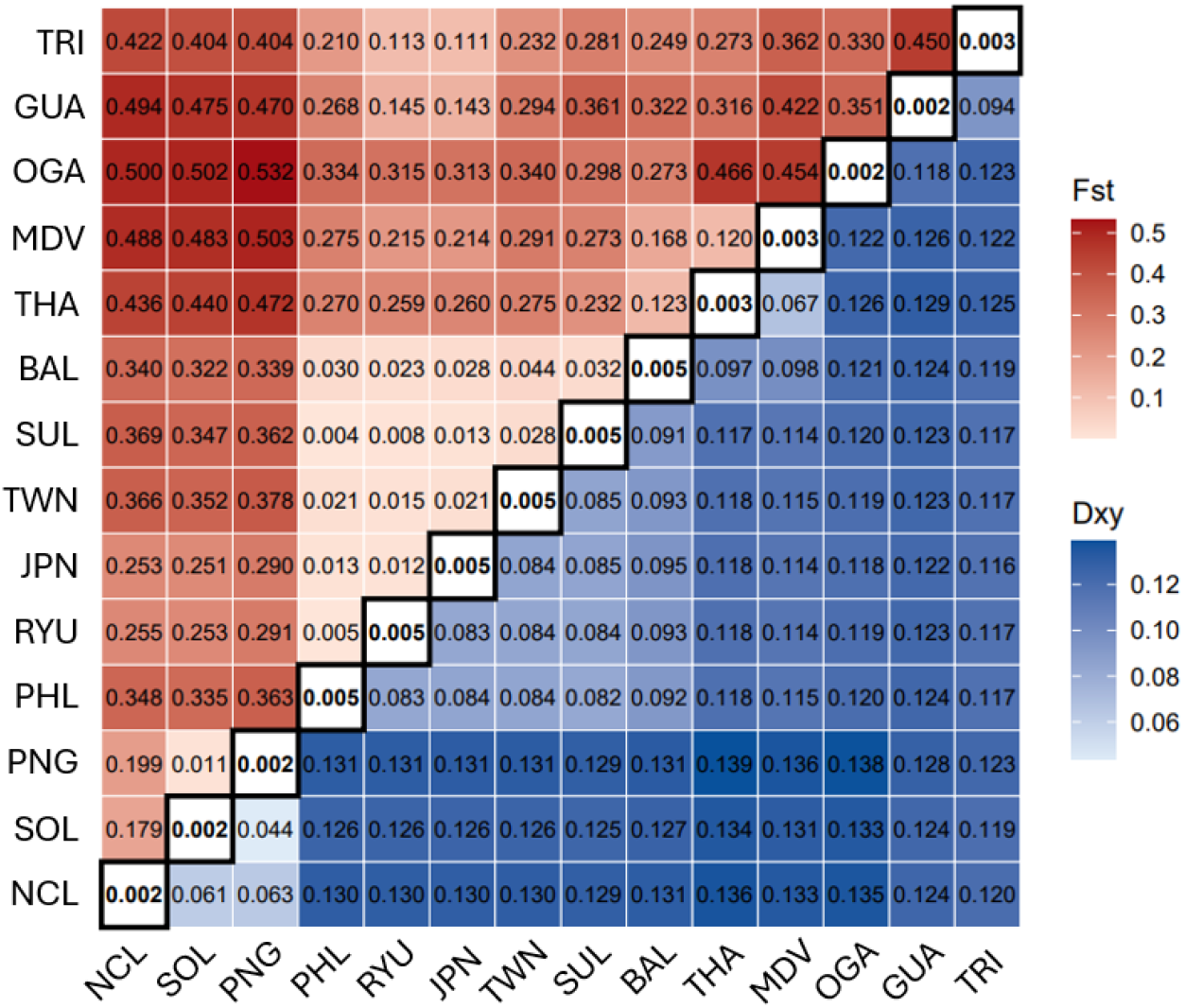
Pairwise population differentiation (*F*_ST_, above diagonal), pairwise population divergence (*d*_xy_, below diagonal) and nucleotide diversity (*π*, diagonal) values for the different populations. Darker colours indicate higher *F*_ST_ and *d*_xy_ values.

To investigate whether some of our samples were related, we ran a kinship analysis, which revealed 36 pairs of individuals with kinship coefficients corresponding to 2^nd^ degree relationships (0.177 – 0.088; Table S2). All were located within four populations: Guam (three pairs), Thailand (17 pairs), PNG (15 pairs) and Solomon Islands (one pair). In other words, in our dataset all individuals from Guam, 8 out of 18 from Thailand, 6 out of 11 in PNG and 2 of 9 individuals from Solomon Islands had at least one 2^nd^ degree relative.

Next, we searched for Runs of Homozygosity (RoH), which can be indicative of inbreeding (via the length of RoH) or past bottlenecks (number of RoH). We called a RoH if a window of at least 300 kb contained only homozygous loci (see Materials and Methods). Guam stood out with 423±20 RoH per individual, almost twice as many as Solomon Islands and PNG with 247±69 and 240±12 RoH, respectively (Table S3). Noteworthy is also Ogasawara, with on average 208±11 RoH per individual. Populations from the central Indo-Pacific region ranged from 13±6 in mainland Japan to 39±16 in Taiwan, while Indian Ocean populations had 62±16 (Thailand) and 65±8 (Maldives) RoH per individual. Finally, *A. tricinctus* individuals had 108±6 RoH on average. The length, indicative of recent inbreeding, ranged from 452±35 and 441±7 in Taiwan and Guam, respectively, to 398±21 in Bali and 400±12 in the Maldives. Taken together, RoH suggest that Guam experienced both recent inbreeding and past bottlenecks, whereas central populations other than Taiwan experienced neither.

### Nuclear phylogeny reveals two major clades

We constructed a phylogenetic tree from a random 5% subset of all SNPs using the TVM+F+ASC+R5 model in IQtree2. The phylogeny revealed that all *A. clarkii* and, notably, *A. tricinctus* samples formed a single monophyletic clade (Fig. 3). Consistent with our PCA and ADMIXTURE results, *A. tricinctus* nested within *A. clarkii* and grouped most closely with the Guam population, although a marked genetic distance separated the two. Two major clades could be observed: One clade containing the Melanesian populations, Ogasawara, Guam and *A. tricinctus*, with Ogasawara splitting off very early, and a clade containing the rest. Most major branches received strong SH-aLRT and bootstrap support (100/100), only for Ogasawara and the Guam & *A. tricinctus* clade it was weaker with 99.5/100 and 97.3/100, respectively.

**Figure 3.**
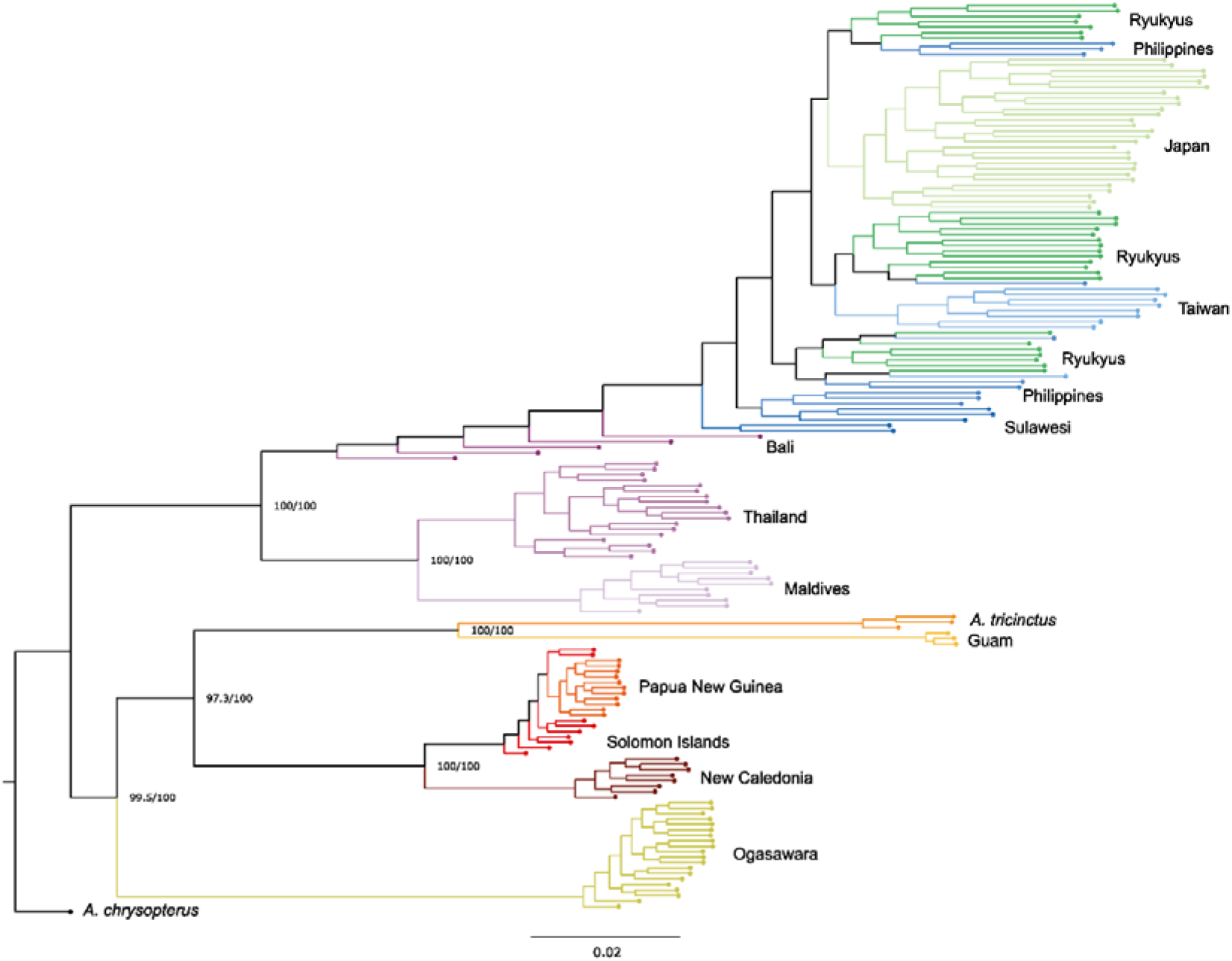
Phylogenetic trees from nuclear DNA. Numbers represent branch support: SH-aLRT/bootstrap values.

We also looked at the phylogenetic structure per chromosome to check for phylogenetic discordance. Pairwise Robinson-Foulds distances (Fig. S4A) revealed that there were only two major organisations of the clades: either Ogasawara branching off early from the Melanesia, Guam and *A. tricinctus* cluster (in 18 chromosomes; Fig. S4B), or branching off early from the Central and Indian Ocean cluster (in 6 chromosomes; Fig. S4C). However, Ogasawara consistently branched off soon after the initial divergence, suggesting that both scenarios did not differ substantially. Additionally, the chromosomal trees revealed several chromosomes (chromosomes 1, 2, 7 and 18) with inverted structures in the Central clade (example in Fig. S4D), however, since this was not linked to the global organisation in *A. clarkii* we did not investigate this further.

Overall, the same groups could be identified as in the PCA and ADMIXTURE analyses. Interestingly, while not perfect, most individuals from the Central population grouped with other individuals from their population.

### Ongoing gene flow from very isolated populations and *A. tricinctus*

We used fastsimcoal2 to simulate the demographic history of our newly sequenced populations: Ogasawara, Guam and *A. tricinctus*. For the Ogasawara population, we tested the phylogenetic topologies (Fig. S4B/C) to confirm whether Ogasawara emerged from the Central or the Melanesian population (Fig. S3 for all tested models). Both, Akaike Information Criterion (AIC = 182,011,517) and the highest likelihood (Fig. 4A) chose the same scenario as the best among the tested ones. In that scenario (Fig. 4B), Ogasawara diverged from the Central population, and there is ongoing gene flow between all the populations. Parameter estimation suggested that Ogasawara diverged much later than the Melanesian populations, 8,863 generations ago, compared to 60,060 generations for Melanesia. Furthermore, the population sizes in Ogasawara (6,080 (haploid sizes)) and Melanesia (9,847) were comparatively small relative to the Central region (44,500; Table 1A). Contemporary gene flow to and from Ogasawara tended to be lower compared to the other comparisons (magnitude E-06 compared to E-05), except gene flow from the Central population to Ogasawara, which was comparable to rates between the Central and Melanesian populations.

**Figure 4.**
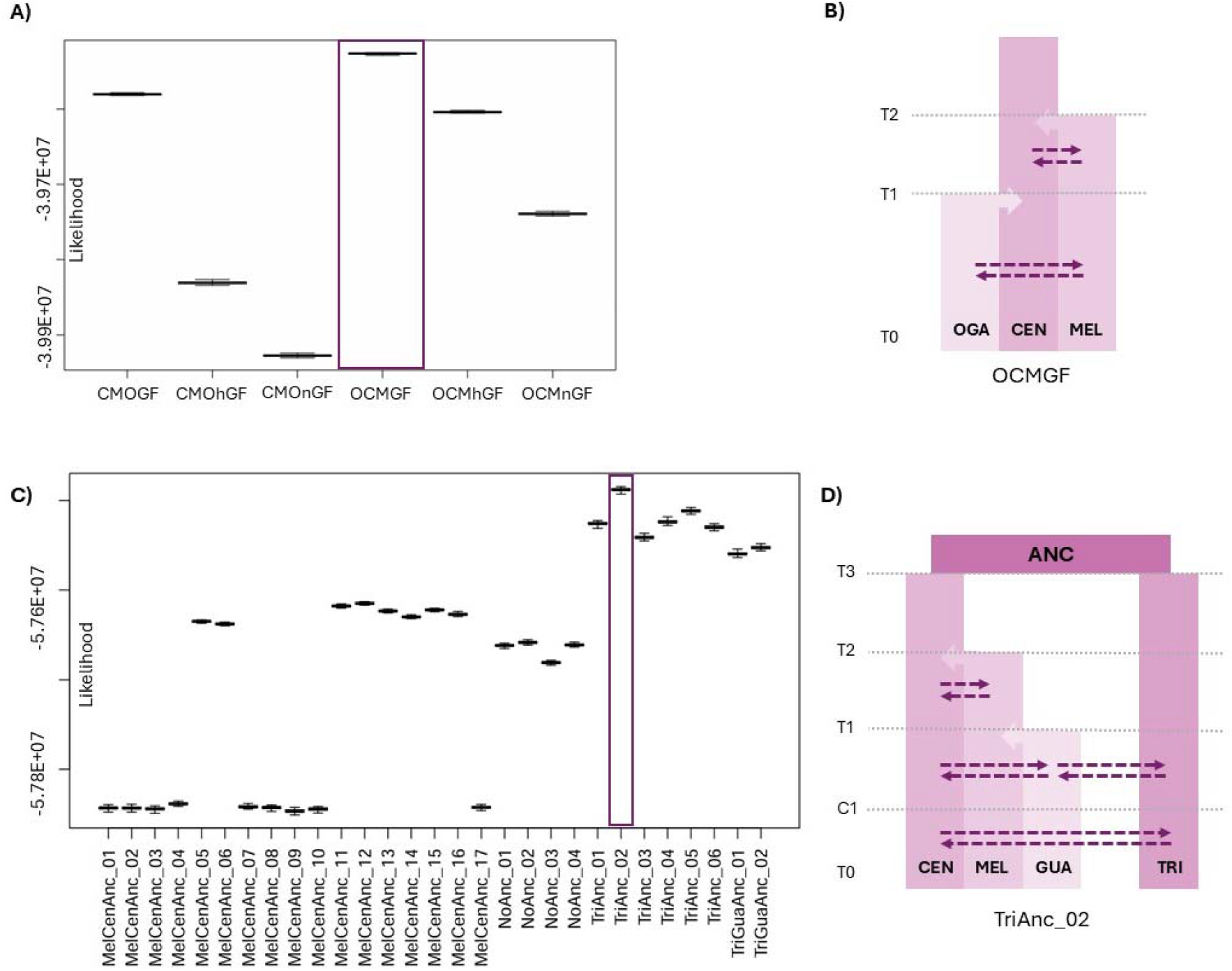
Best demographic scenarios and model likelihoods from fastsimcoal2 simulations. **A, B)** Likelihoods of tested model and best scenario for Ogasawara, respectively. **C, D)** Likelihoods of tested models and best scenario for Guam and *A. tricinctus*, respectively. Dashed arrows represent geneflow between any of the overlapping populations. Light arrows depict merging of populations (fastsimcoal2 simulations go back in time). T0 = current time; T1 = split time from Ogasawara and Guam, respectively; T2 = Split time from Melanesia (defined as 60,060 generations); T3 = Split time of ancestral population into *A. clarkii* and *A. tricinctus*; C1 = time of first contact between *A. tricinctus* and *A. clarkii*.

**Table 1:**
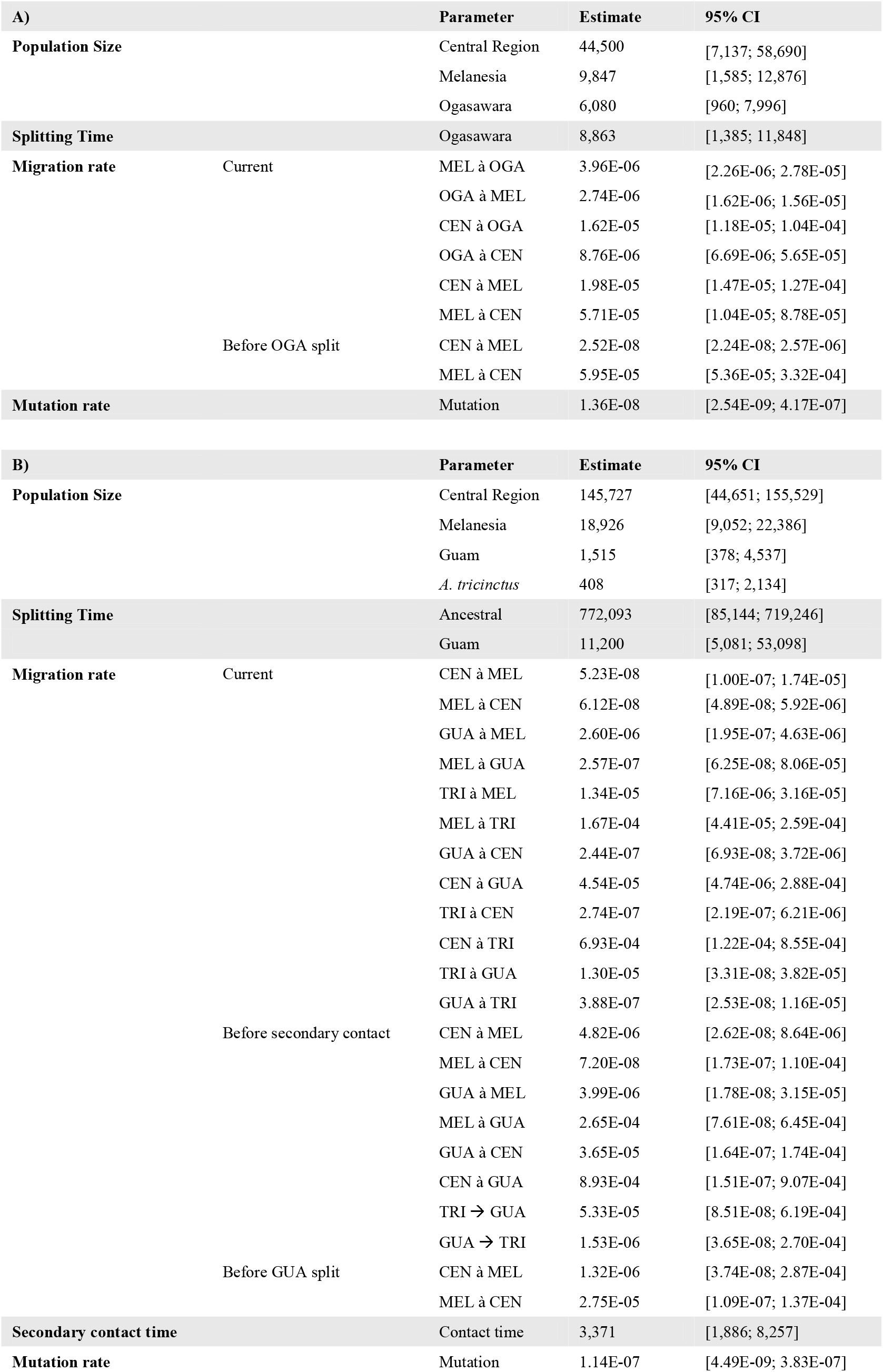
Parameter estimates of the best scenarios from fastsimcoal2 modelling. **A)** Ogasawara splitting from the Central population. **B)** Best model for Guam and *A. tricinctus*; *A. clarkii* and *A. tricinctus* diverging from an ancestral population, with more recent secondary contact between the rest. Abbreviations: CI confidence interval; MEL Melanesia; OGA Ogasawara, CEN Central population; GUA Guam; TRI *A. tricinctus*.

In the scenario involving Guam and *A. tricinctus,* the best model (AIC = 264,724,812; highest likelihood; Fig. 4C) suggested an early separation between *A. clarkii* and *A. tricinctus*, with an initial secondary contact with Guam, and later with all populations (Fig. 4D, Table 1B). Estimated population sizes for the Central and Melanesian populations were 145,727 and 18,926, respectively, much larger than the ones in the Ogasawara simulations. Guam and *A. tricinctus* were the smallest, with only 1,515 and 408, respectively (haploid sizes). The highest gene flow was suggested from Central to *A. tricinctus* and Melanesia to *A. tricinctus*, with the lowest in both directions between Central and Melanesia, however, the confidence intervals for several migration rates were wide, sometimes covering three orders of magnitude, and some rates falling outside the confidence intervals (e.g. Central to Melanesia).

Expanding our fastsimcoal2 simulations to include more distantly related clades, we calculated Dsuite’s f-branch statistics (*f_b_*) which can highlight phylogenetic branches with excessive allele sharing, an indication for introgression. Our analysis revealed introgression between the ancestral branch from the Guam and *A. tricinctus* clade and the current Central and Indian Ocean clade (Fig. 5). This was particularly driven by *A. tricinctus*, which showed signs of introgression with all tested populations but Ogasawara. The latter population, on the other hand, showed pronounced introgression with Guam (*f_b_* = 0.103). As supported by the mitochondrial and nuclear trees, Bali, for this analysis located in the Central clade, showed strong introgression with Indian Ocean populations (*f_b_* = 0.239 – 0.300).

**Figure 5.**
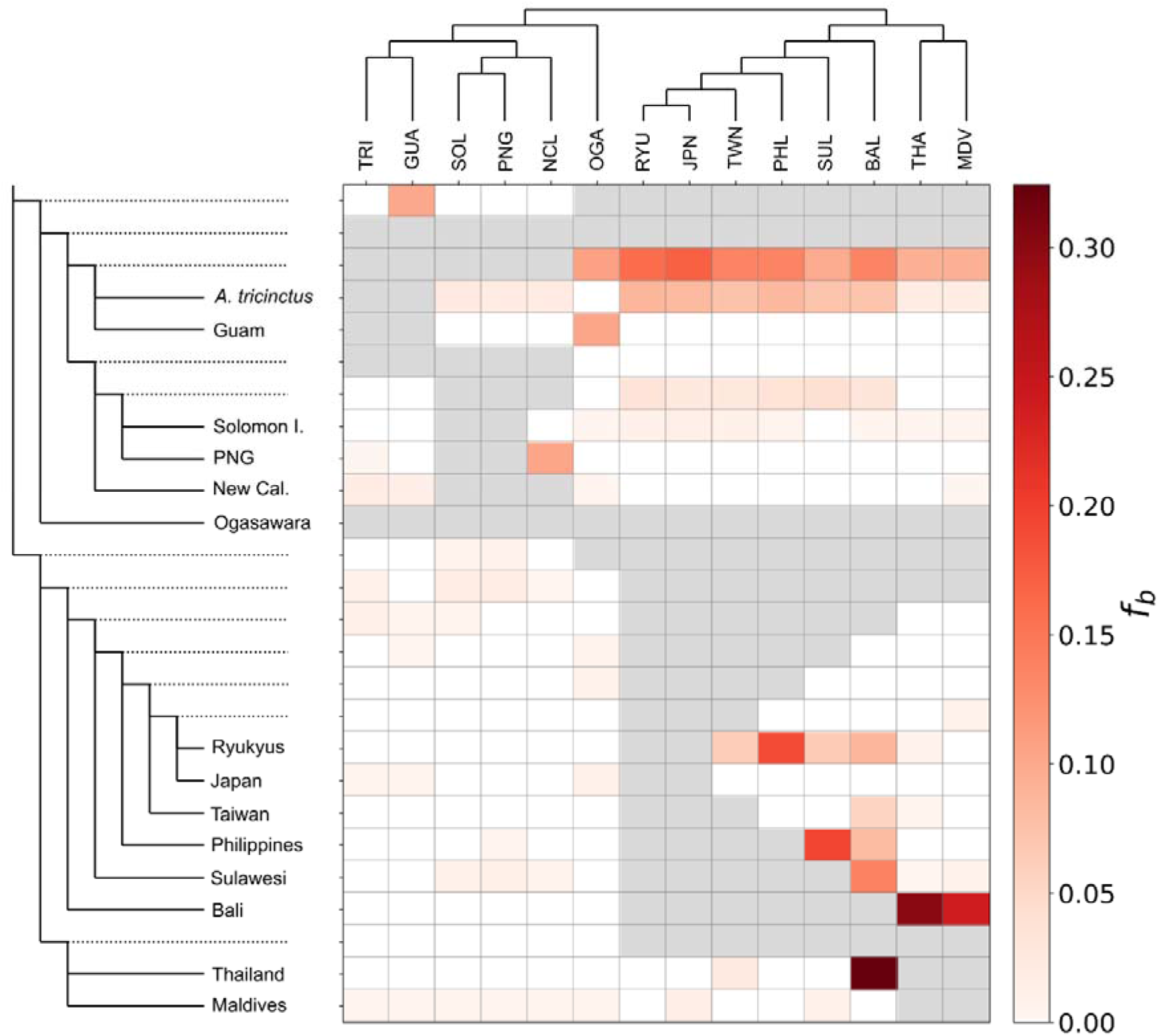
Visualisation of fbranch statistic (*f_b_*), showing branches with excessive allele sharing in red. Dotted lines represent ancestral nodes. Combinations that could not be computed due to the tree layout are shown in grey.

### Mitochondrial phylogeny shows deep divergence within *A. clarkii*

To put the divergence within *A. clarkii* into perspective with other anemonefish species, we also made a phylogenetic tree using mtDNA and added mitochondrial sequences from other *Amphiprion* species pairs: *A. frenatus, A. ephippium, A. perideraion, A. akallopisos*, *A. percula,* and *A. ocellaris*. The mitochondrial phylogeny (Fig. 6) revealed a very strong separation between *A. clarkii* from Melanesia and *A. clarkii* from the other regions. The previous major divisions (Indian Ocean, *A. tricinctus*, Guam, and Ogasawara) were still distinct. One external *A. clarkii* from the Paracel Islands (South China Sea; NC_023967.1) also clustered with the Central population. Of note, the cophenetic distance between *A. clarkii* from Melanesia and elsewhere (0.049) was greater than the distance between the other species pairs included: 0.027 for *A. perideraion / A. akallopisos,* 0.013 for *A. frenatus / A. ephippium*, and 0.047 between *A. percula* and *A. ocellaris*.

**Figure 6.**
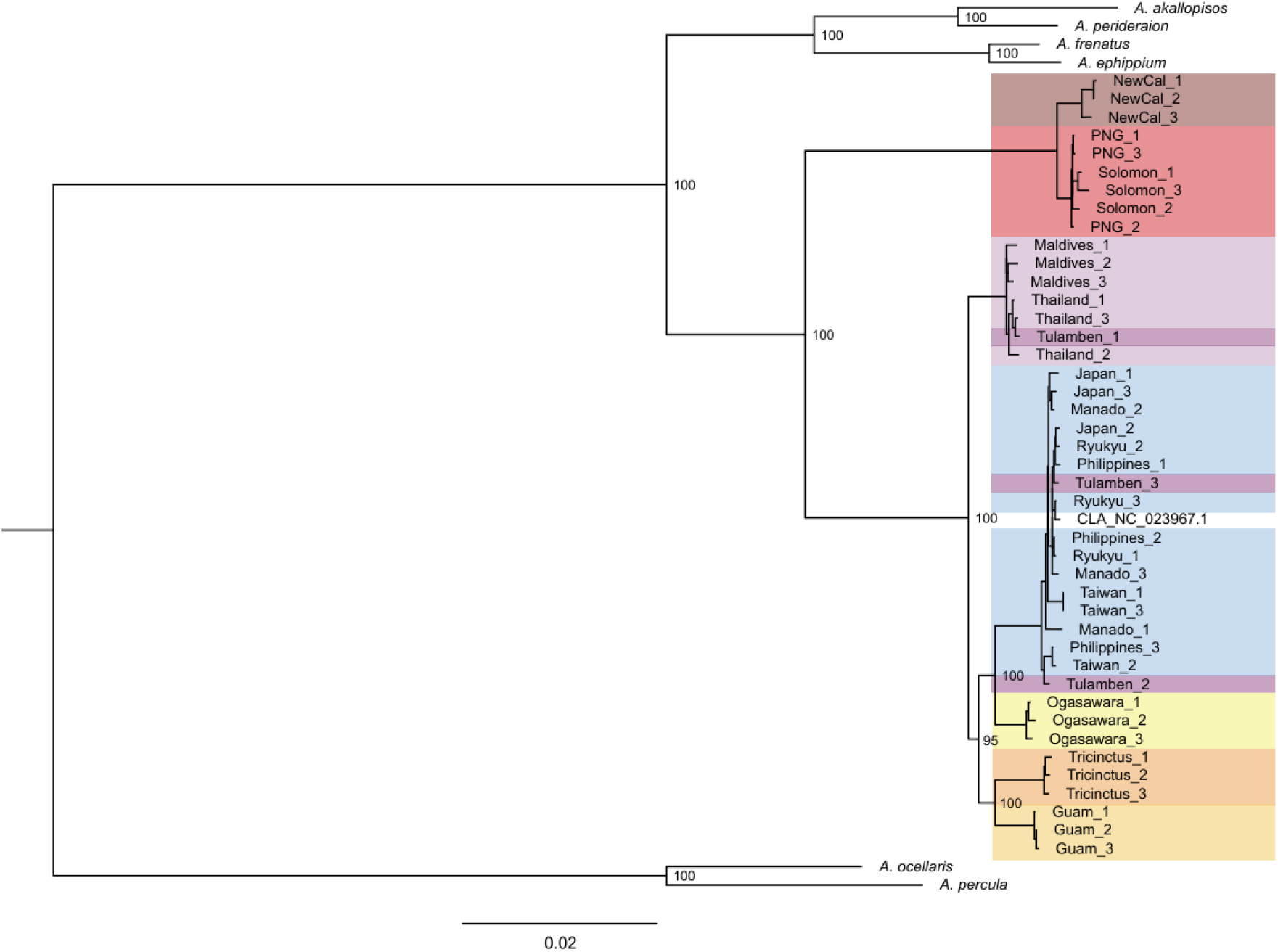
Mitochondrial tree of three random individuals per population, based on the coding sequences of the mitochondrial DNA. Other species and a publicly available *A. clarkii* individual (CLA_NC_023967.3) included to have a wider comparison. Node numbers represent the ultrafast bootstrap values. The tree was visualised with figtree v1.4.4.

## Discussion

With its wide distribution that includes the centre of biodiversity, but also remote archipelagos such as Ogasawara, and its host generalism, *A. clarkii* is an intriguing anemonefish species to study population genomic characteristics in both, well-connected and isolated populations. Here, we compiled the most complete set of whole genome data for *A. clarkii*, and for any *Amphiprion* species. As an expansion of Schmid et al. (2024), we showed that the well-connected central Indo-Pacific population extended to the northern range limit of this species, however, other major clades were strongly diverged. Its close relative *A. tricinctus* was consistently placed within *A. clarkii,* forming a clade with *A. clarkii* from Guam, suggesting that the current taxonomy may not reflect the real situation.

### Five clades form the *A. clarkii* species complex

Through all our analyses, five main clades emerged for this species complex: Central Indo-Pacific (from Indonesia to Japan), Indian Ocean (Thailand and Maldives), Melanesia (Solomon Islands, Papua New Guinea and New Caledonia), and Ogasawara lineages, and the Guam lineage combined with *A. tricinctus*. The Central, Indian Ocean and partially the Melanesian populations had low internal *F*_ST_ values, indicating high internal connectivity and low structure. By adding recent samples from Thailand (García Jiménez et al., 2025) to our dataset, we confirmed that at least parts of the Indian Ocean also form a well-connected basin with low population structure. The Central population, acting as a genetic hub and previously established as the species’ origin (Schmid et al., 2024), had no indications of reduced diversity or increased isolation at the northern range limit in Japan.

We observed a clear reduction in genetic diversity in all the clades outside the Central region. Interestingly, we found signatures of transition or secondary contact between the Central and Indian Ocean populations in the Bali samples. In the nuclear genome analyses, these individuals were closer to the Central population but branched off one by one shortly after the major separation between Indian Ocean and the Central group. In contrast, in the mitochondrial genome, the Bali individuals were either completely within the Indian Ocean or Central group. Secondary contact was also suggested by the fbranch statistics, with high values between Bali and the Indian Ocean groups. In addition to Schmid et al. (2024), similar results were also reported by Ducret et al. (2022), who showed Bali and Javan specimens consisting of a mix between Central and Indian Ocean groups. A mosaic of connectivity and isolation, as well as contact zones, as observed here in *A. clarkii*, are common in widespread reef fishes, for instance in the widespread grouper *Cephalopholis argus* (Gaither et al., 2011). Our large genomic dataset included individuals of representative populations from almost the entire *A. clarkii* range. The main remaining geographic gap is the western range edge in the Persian/Arabian Gulf. Nevertheless, the high connectivity reported between western and eastern Indian Ocean populations of *A. akallopisos* (Marcionetti et al., 2024) suggests that long-distance connectivity across the entire basin can also be expected in *A. clarkii*.

### Signs of inbreeding and historic bottlenecks in some populations

Our kinship analysis revealed 2^nd^ degree relationships between all the samples from Guam. Despite great sampling efforts we could only obtain sequences from three individuals from two sites. Even though anemonefishes have a two-week oceanic larval phase (Jones et al., 2022), they often tend to settle in their natal area, as shown with *A. percula* from Kimbe bay where many relatives could be found (Salles et al., 2016). Nevertheless, the chances of only sampling 2^nd^ degree relatives are rather small and indicate a population with potential inbreeding.

High number of shorter Runs of Homozygosity (RoH) indicate historic bottlenecks, whereas longer RoH are a sign of recent inbreeding (Ceballos et al., 2018; Kirin et al., 2010). Our data showed how individuals from Taiwan had the longest segments, suggesting potential inbreeding, although the difference in RoH lengths with the shortest populations was only around 10%. In contrast, the number of RoH varied greatly, with individuals from Guam having almost twice as many RoH than individuals from Solomon Islands, Papua New Guinea and Ogasawara, other populations with many RoH. Since each individual was searched for RoH separately, the relatedness of the Guam samples should not inflate this count. Instead, it is a strong indication of a severe bottleneck in the past and could be a result from founder effect (Jensen et al., 2021). Interestingly, *A. tricinctus* individuals had fewer RoH than individuals in other isolated populations, corresponding to a weaker bottleneck signature. While not examining RoH directly, a study on the grouper *Cephalopholis argus* found populations in the Marshall Islands (home to *A. tricinctus*) to be more admixed and connected than those in Guam (Amirthalingam et al., 2025), potentially explaining the weaker bottleneck signature in *A. tricinctus* relative to other isolated populations.

### Phylogenies show strong intraspecific divergence

Through all the analyses, the degree of divergence we observed among *Amphiprion clarkii* populations was remarkable and, in several cases, equivalent to that observed between recognised *Amphiprion* species. Particularly in the mitochondrial tree, the Melanesian populations (Papua New Guinea, Solomon Islands, New Caledonia) were separated from all others by levels of differentiation similar or greater to those between *A. percula* and *A. ocellaris, A. frenatus* and *A. ephippium,* and *A. perideraion* and *A. akallopisos.* Branches that are longer within the *A. clarkii* complex compared to other species pairs in a single tree can also be observed in other phylogenies, both in trees from nuclear (Gaboriau et al., 2025) and mitochondrial DNA (Litsios et al., 2014b; Nguyen et al., 2020).

There were some noticeable differences between our nuclear and mitochondrial trees: While the mitochondrial genome suggested that the Melanesian population split off very early, and were therefore by far the most diverged clade, the nuclear tree suggests that *A. tricinctus* and Guam, as well as Ogasawara in most chromosomes, branched from the Melanesian rather than the Central cluster. Such differences between mitochondrial and nuclear phylogeny are common and known as mito-nuclear discordance (reviewed in Toews & Brelsford, 2012). The mito-nuclear discordance has been attributed to, among others, mitochondrial introgression and incomplete lineage sorting (Bonnet et al., 2017; Degnan & Rosenberg, 2009; Toews & Brelsford, 2012). Given the observed gene flow between populations, mitochondrial introgression followed by a drift-driven fixation in small and isolated populations may explain the observed discordance. Such mitochondrial introgression has also been described in the skunk complex (*A. akallopisos* group; Marcionetti et al., 2024) and are therefore not uncommon in this genus.

### Simulations and introgression tests indicate gene flow even among isolated populations

In both tested population groups (Central, Melanesia and either Ogasawara, or Guam and *A. tricinctus*) our demographic simulations showed ongoing gene flow between all populations, consistent with Schmid et al. (2024). For Ogasawara, Ogasawara splitting from the Central population was the most likely scenario, and while this was the branching supported by fewer chromosomal phylogenies (Fig. S4C) and contradicting the whole-genome tree, it followed the mitochondrial tree structure. The Ogasawara population was also the smallest, although its size difference to Melanesia was smaller than expected based on limited habitat areas and relatively low abundance, at least around Chichi-jima (JZ, JDR, MM, personal observation). Population structure can affect estimations of effective population sizes (Charlesworth, 2009), providing a potential explanation for the probable overestimation of the Ogasawara *A. clarkii* effective population size. Due to the remoteness of Ogasawara, it is no surprise that migration rates to and from Ogasawara tended to be the lowest, indicating a high level of isolation of the local *A. clarkii* population. Similar isolation effects in Ogasawara have been observed in the reef-building coral *Galaxea fascicularis* (Wepfer et al., 2022).

For the models involving Guam and *A. tricinctus*, the best model among the tested ones was with an ancestral population that subsequently split into *A. clarkii* and *A. tricinctus*, with a more recent secondary contact. It has been noted that model selection tends to favour a secondary contact scenario if inadequate demographic history is provided, especially regarding population size history (Momigliano et al., 2021; Smith & Hahn, 2024). Following common practice (reviewed in Momigliano et al., 2021), we simplified our models by not including changes in ancestral population sizes to not inflate the number of parameters to be simulated. Instead, we tested an ancestral population splitting into Melanesia and Central populations, to give various secondary contact scenarios. However, their likelihood was lower. The best of our tested scenarios suggested *A. clarkii* and *A. tricinctus* having two separate historical trajectories with more recent secondary contact, first with Guam and then with the remaining populations. This close contact with Guam would explain the current situation despite separate histories. However, it contradicts the expectation, based on Schmid et al. (2024), of a stepping-stone model without a shared ancestral population, which was also supported with our simulations of Central and Melanesian populations not emerging from a shared ancestral population, despite being the least-connected. Whether the small sample sizes for Guam and *A. tricinctus* were a driver of these results cannot be ruled out, however, for all our population genetic analyses the results were consistent between the full data set and a reduced dataset with three individuals per population.

Furthermore, we observed wide confidence intervals, some with three orders of magnitude (E-08 to E-05), and a few parameters outside the CI, indicating pronounced uncertainties in the parameter estimation, mainly in estimates involving Guam. Wide CI are not uncommon in fastsimcoal2 simulations (De Jong et al., 2023; Ruffley et al., 2018; Tengstedt et al., 2025), as are parameters beyond the CI (e.g. Postaire et al., 2024; Tengstedt et al., 2025), and are a reminder that all the models are highly simplified.

Previously, Schmid et al. (2024) showed that despite the strong divergence, gene flow is ongoing between the central, Indian Ocean and Melanesian populations. Here, the results from demographic modelling confirmed ongoing gene flow between the Central and Melanesian populations, as well as between them and isolated *A. clarkii* from Ogasawara and Guam and *A. tricinctus*. While we decided to not run a global model due to the vast number of possible splitting orders, we could close this gap with f-branch statistics (*f_b_*). Using Dsuite, we confirmed gene flow, or introgression, between populations not directly compared in fastsimcoal2 simulation, for instance Guam and Ogasawara.

### Taxonomic implications of strong divergence and *A. tricinctus* position

How to define a species remains one of the most debated questions in evolutionary biology, and different species concepts, such as biological, phylogenetic or evolutionary, often lead to different conclusions (De Queiroz, 2007; Mayr, 1996), thus affecting species delimitation. The strong divergence of different populations, and the inclusion of *A. tricinctus* within *A. clarkii*, yet with ongoing gene flow between all populations, make this clade an interesting example of challenging species delimitation.

The taxonomic history of *A. clarkii* is complex. After decades of taxonomic reorganisation, *A. clarkii* has nine synonyms, including *Amphiprion papuensis* Macleay, 1883 for New Guinea and Queensland and *A. snyderii* Ishikawa, 1903 for Ogasawara. *Amphiprion clarkii* (Bennett, 1830) itself was initially described from a Sri Lankan specimen, and is thus from the Indian Ocean, whereas the first description of a species from the Central population of this study was *Amphiprion japonicus* Temminck & Schlegel, 1843, from a Japanese specimen.

The lack of clarity in *A. clarkii* phylogenetics is not unique among anemonefishes, as shown by the recent description of *A. maohiensis* (O’Donnell et al., 2025) and the ongoing discussion as to whether *A. polymnus* is a single polymorphic species or a complex of multiple species (Fitzgerald et al., 2026). *A. maohiensis* is not within the *A. chrysopterus* complex based on genomic divergence, both nuclear and mitochondrial, allopatry and subtle morphological differences, dorsal fin length and peduncle colour (O’Donnell et al., 2025). Similarly, Fitzgerald et al. (2026) observed strong genomic divergence, as well as distinct pigmentation patterns between the different groups of *A. polymnus*, yet they avoided a clear conclusion about the status of that species complex. In the current study, we were limited to genomic data and lacked the phenotypic components seen in other studies (O’Donnell et al., 2025; Fitzgerald et al., 2026), making a formal taxonomic conclusion unadvisable. However, whether phenotypic data would solve the issues of *A. clarkii* delimitation remains doubtful. Even within Japan, Bell et al. (1982) noted remarkable differences in numbers of dorsal spines and pectoral rays, as well as distinct tail colourations, for different populations. Furthermore, they described a northward increase in melanisation, which was recently confirmed quantitively (Mercader et al., in prep.). Of additional challenge is the effect of host anemone on colour morphs; black morphs in *Stichodactyla* and orange for those associating with *Entacmaea, Heteractis* and *Radianthus* (Allen, 1972, Militz et al., 2016). However, colour assessment from the literature is difficult as often the host anemone species is not included in reports and pictures, and the confirmation of geographic effects on colouration ultimately requires comparisons separated by host species. In addition to morphology, a best practice approach would also include behaviour, reproduction, and ecological traits (De Queiroz, 2007, Petzold & Hassanin, 2020). In anemonefish, all sharing the same ecological niche (Allen, 1972) and have been found to naturally hybridise (Ollerton et al., 2007; Schmid et al., 2025), finding such distinct characteristics can be challenging, further increasing the complexity of future species delimitation. However, we believe that a careful morphological analysis of *A. clarkii* using microCT scans with samples coming from the whole geographical range could bring valuable characters to eventually confirm or dismiss a species status for some of the populations we discuss in this paper.

The strong separation of the Melanesian, Guam and Ogasawara *A. clarkii* lineages may correspond to independent evolutionary trajectories that have reached the stage of reproductive autonomy. The persistence of gene flow throughout the evolutionary history of this complex, however, prevents a straightforward interpretation of these divergent lineages as distinct species (Schmid et al., 2024). While the absence of gene flow is a common species criterion (Mayr, 1996; Westram et al., 2022), recent studies show that speciation with gene flow is not uncommon (Capblancq et al., 2019; Wang et al., 2016). In such cases, only a limited portion of the genome, typically regions under divergent selection, may remain isolated while the rest of the genome continues to exchange information (Cadena & Zapata, 2021; Vaux et al., 2021). In the end, splitting or lumping in such cases depend on what criteria are to be applied for species delimitation, and whether researchers allow for limited gene flow between potential species.

Our data strongly confirm the previous indications that *A. tricinctus* is positioned within the *A. clarkii* clade (Litsios & Salamin, 2014). Even though our demographic modelling suggests a secondary contact scenario, all the other analyses firmly place *A. tricinctus* within *A. clarkii*, and gene flow between *A. tricinctus* and some of the *A. clarkii* clades was observed to be stronger than between different *A. clarkii* populations. Therefore, this strongly suggests the need for taxonomic revision of the *A. clarkii* species complex. According to the original description of *A. tricinctus,* its main difference with *A. clarkii* is the body depth in adults, black caudal fins, and the second bar width (Allen, 1972; Schultz, 1953), along with the lack of range overlap. However, due to the great variability of *A. clarkii,* none of these characteristics is unique to *A. tricinctus*, as even *A. clarkii* juveniles can have black caudal fins. That said, there are two possibilities on how to resolve the *A. clarkii* conundrum: either splitting the *A. clarkii* species complex into different species; or reevaluating the taxonomic rank of *A. tricinctus* to integrate it into *A. clarkii*. However, keeping *A. tricinctus* as an independent species without splitting *A. clarkii* would clearly contradict our findings.

Interestingly, Whitley (1929), who re-described *A. papuensis*, predicted that similar species would be later grouped together, and that a subsequent need for re-separation could be necessary. Except *A. clarkii* from Guam, all our major clades have been previously described as independent species, and therefore Whitley’s prediction may ultimately prove true.

## Acknowledgements

We thank Masanori Nakae and the Fish Section and Center for Molecular Biodiversity Research of the National Museum of Nature and Science, Japan for providing *A. clarkii* samples from Kochi and Daito, and Ken Maeda for facilitating the logistics. We also thank Hiroki Takamiyagi, Erina Kawai, Jeff Jolly, Timothy Ravasi (all OIST), Emmeline Jamodiong (U. Ryukyus), Sylvain Agostini (Shimoda Station, Tsukuba U.), Daisuke Uyeno (Kagoshima U.) and Midori Matsuoka (Kagoshima U.) for fieldwork assistance in Japan, and Héctor Torrado and Thomas Payne and University of Guam for the sample collection in Guam, as well as all the local authorities. We thank Shu-Hua Li for obtaining *A. tricinctus* fin clips from the breeding pair at Academia Sinica, Taiwan. Furthermore, we express our gratitude to Marcela Herrera, Emma Gairin, Yi Ming Weng, Sarah Schmid, Alberto García Jiménez and Wan-Ting Huang for discussions and guidance with analyses, and Lucy Fitzgerald and Floriane Coulmance for critical reading of the manuscript. We also thank the Marine Science, Sequencing, and Scientific Computing and Data Analysis sections at the Okinawa Institute of Science and Technology for their assistance.

## Author contributions

JZ, MM and VL designed research. JZ, MM, CL and JDR contributed to sample collection. JZ, AM and SM performed research, lab work and data analysis. JZ wrote the manuscript. All authors read, commented and approved the final version of the manuscript.

## Data availability statement

The sequences are deposited in the SRA (BioProject PRJNA1524052). The scripts used for data analysis are available on https://github.com/Jann-Z/Clarkii_complex_WGS and will be posted on Dryad when accepted.

## Funding statement

This work was supported by an OIST KICKS Program for Research Scholarship Fund and JSPS DC2 scholarship (grant no. 25KJ2246). J.D.R. gratefully acknowledges the Japan Society of the Promotion of Science (JSPS) International B research scheme project ‘Marine environmental DNA metabarcoding: understanding oceanic islands and mainland sites across anthropogenic gradients in two hemispheres’ (grant no. 20KK0164).

## Conflict of interest statement

The authors have no conflict of interest to declare for the presented work.

## Ethics approval statement

Sampling was carried out according to local laws and regulations and with permission of local fishery unions. Special permits were obtained for protected areas: Aka (Kerama Shoto National Park, Japan: Permit Number 2106291), Ogasawara (Permit Number 4-48), and Guam (License Number SCR-24-007)

## Notes

### Competing Interest Statement

The authors have declared no competing interest.

