## Supplemental Figures and Tables for "One species or several? Genomic evidence for deep divergence within the widely distributed anemonefish *Amphiprion clarkii* species complex"

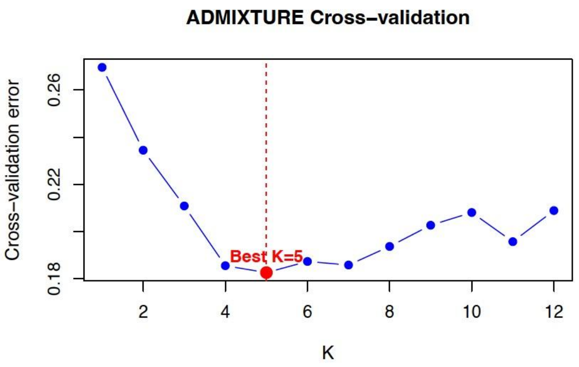


**Figure S1:** Cross validation errors for *K=1-12* from ADMIXTURE.

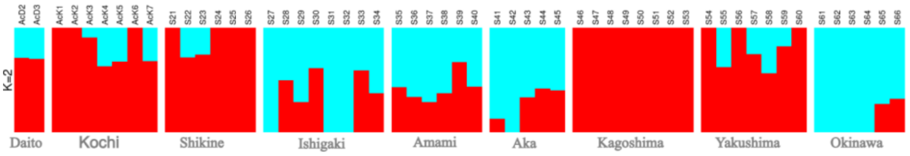


**Figure S2:** Admixture result for samples from Japan (without Ogasawara) for *K = 2*. Ishigaki, Amami, Aka, Okinawa, Daito are part of the Ryukyus; Kochi, Shikine, Kagoshima are considered mainland Japan, Yakushima is close to Kagoshima.

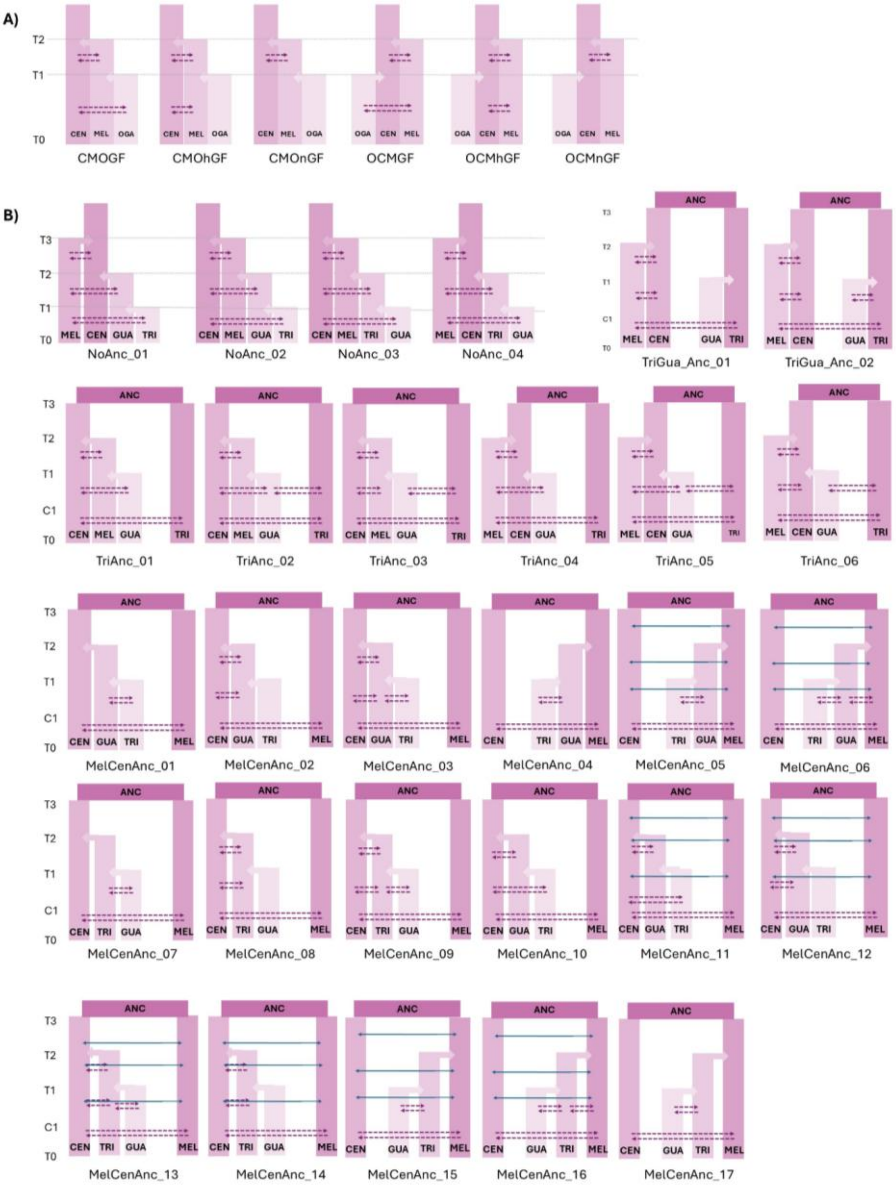


**Figure S3:** The models tested for fastsimcoal2 simulations. **A)** Models involving Ogasawara; **B)** models involving Guam and *A. tricinctus.* Dashed arrows indicate gene flow between all the covered populations; solid arrows indicate gene flow only between the two end points (CEN and MEL). CEN Central; MEL Melanesia; OGA Ogasawara; GUA Guam; TRI *A. tricinctus.*

**Commented [AM1]:** Check the figure of the best model reported in the main text as it is not the same as here.

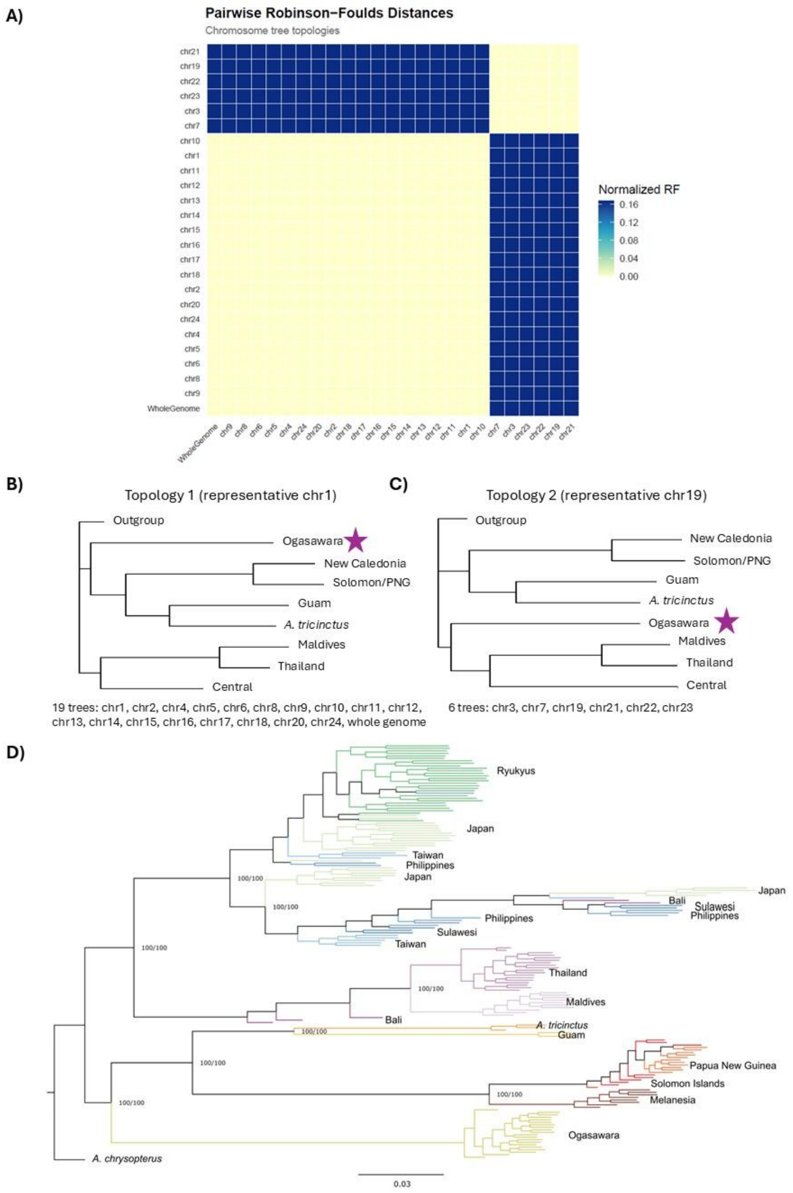


**Figure S4:** Phylogenetic tree topologies. **A)** Pairwise Robinson-Foulds Distances between each chromosome and the whole genome. Colours indicate distances. **B)** Topology 1, representing 18 chromosomes and the whole genome. **C)** Topology 2, representing 6 chromosomes. **D)** Example of a tree with inversion structures in the central population (chromosome 1). Ogasawara in the topologies is marked with a purple star.

**Table S1.** Sample information for all 166 samples used in this study. All new samples are deposited in the SRA under BioProject PRJNA1524052.

| Sample name | Location | Population | Collection | Sequencing depth (mean) | Reads |
|---|---|---|---|---|---|
| AcD2 | Minamidaito | Ryukyu | National Museum of Nature and Science, Japan | 28.9 | 206,506,726 |
| AcD3 | Minamidaito | Ryukyu | National Museum of Nature and Science, Japan | 27.1 | 204,720,714 |
| AcK1 | Okinoshima | Japan | National Museum of Nature and Science, Japan | 24.7 | 178,001,666 |
| AcK2 | Okinoshima | Japan | National Museum of Nature and Science, Japan | 27.7 | 193,973,762 |
| AcK3 | Okinoshima | Japan | National Museum of Nature and Science, Japan | 26.8 | 190,885,982 |
| AcK4 | Okinoshima | Japan | National Museum of Nature and Science, Japan | 27.6 | 204,396,792 |
| AcK5 | Okinoshima | Japan | National Museum of Nature and Science, Japan | 24.3 | 172,539,328 |
| AcK6 | Okinoshima | Japan | National Museum of Nature and Science, Japan | 20.8 | 146,164,628 |
| AcK7 | Okinoshima | Japan | Museum of Nature and Science, Japan | 20.3 | 143,416,198 |
| At1 | Captivity | *A. tricinctus* | Breeding couple at Academia Sinica | 25.2 | 183,157,580 |
| At2 | Captivity | *A. tricinctus* | Breeding couple at Academia Sinica | 25.8 | 185,501,692 |
| CLK_113 | Phi Phi Islands | Thailand | PRJNA1202426 [1] | 20.1 | 197,387,472 |
| CLK_120 | Phi Phi Islands | Thailand | PRJNA1202426 [1] | 20.6 | 207,738,304 |
| CLK_123 | Phi Phi Islands | Thailand | PRJNA1202426 [1] | 18.8 | 177,642,860 |
| CLK_130 | Phi Phi Islands | Thailand | PRJNA1202426 [1] | 17.9 | 169,146,970 |
| CLK_133 | Phi Phi Islands | Thailand | PRJNA1202426 [1] | 21.3 | 213,761,342 |
| CLK_150 | Ko Lanta | Thailand | PRJNA1202426 [1] | 15.5 | 151,242,984 |
| CLK_158 | Ko Lanta | Thailand | PRJNA1202426 [1] | 18.8 | 185,339,406 |
| CLK_161 | Ko Lanta | Thailand | PRJNA1202426 [1] | 14.9 | 147,726,128 |
| CLK_166 | Ko Lanta | Thailand | PRJNA1202426 [1] | 18.2 | 168,038,766 |
| CLK_169 | Ko Lanta | Thailand | PRJNA1202426 [1] | 14.1 | 128,208,676 |
| CLK_65 | Phuket | Thailand | PRJNA1202426 [1] | 17.3 | 162,421,400 |
| CLK_68 | Phuket | Thailand | PRJNA1202426 [1] | 19.1 | 190,491,531 |
| CLK_71 | Phuket | Thailand | PRJNA1202426 [1] | 17.2 | 158,533,500 |
| CLK_76 | Phuket | Thailand | PRJNA1202426 [1] | 18.8 | 177,922,800 |
| CLK_85 | Similan Islands | Thailand | PRJNA1202426 [1] | 17.8 | 165,453,717 |
| CLK_90 | Similan Islands | Thailand | PRJNA1202426 [1] | 19.1 | 177,830,985 |
| CLK_91 | Similan Islands | Thailand | PRJNA1202426 [1] | 22.4 | 216,721,000 |
| CLK_95 | Similan Islands | Thailand | PRJNA1202426 [1] | 20.5 | 201,885,000 |
| Ac76 | Ogasawara | Ogasawara | Own | 15.1 | 140,865,368 |
| Ac77 | Ogasawara | Ogasawara | Own | 15.0 | 140,711,020 |

**Commented [AM2]:** I guess that for these saples here, you can cite Alberto's paper (https://onlinelibrary.wiley.com/doi/full/10.1111/mec.70134) and/or the SRA (apparently PRJNA1202426, according to his paper)

| Sample name | Location | Population | Collection | Sequencing depth (mean) | Reads |
|---|---|---|---|---|---|
| Ac78 | Ogasawara | Ogasawara | Own | 14.8 | 138,746,266 |
| Ac79 | Ogasawara | Ogasawara | Own | 14.7 | 137,287,960 |
| Ac80 | Ogasawara | Ogasawara | Own | 13.6 | 124,732,182 |
| Ac81 | Ogasawara | Ogasawara | Own | 15.5 | 147,844,990 |
| Ac82 | Ogasawara | Ogasawara | Own | 18.0 | 172,891,240 |
| Ac83 | Ogasawara | Ogasawara | Own | 13.1 | 120,701,290 |
| Ac84 | Ogasawara | Ogasawara | Own | 16.5 | 158,304,390 |
| Ac85 | Ogasawara | Ogasawara | Own | 14.5 | 134,052,344 |
| Ac86 | Ogasawara | Ogasawara | Own | 16.0 | 151,639,696 |
| Ac87 | Ogasawara | Ogasawara | Own | 15.9 | 150,843,634 |
| Ac88 | Ogasawara | Ogasawara | Own | 16.7 | 156,815,158 |
| Ac89 | Ogasawara | Ogasawara | Own | 17.6 | 170,177,798 |
| Ac90 | Ogasawara | Ogasawara | Own | 15.2 | 144,297,574 |
| Ac91 | Ogasawara | Ogasawara | Own | 17.6 | 168,264,200 |
| Ac92 | Ogasawara | Ogasawara | Own | 17.1 | 162,522,482 |
| Ac93 | Ogasawara | Ogasawara | Own | 15.1 | 143,557,824 |
| Ac94 | Ogasawara | Ogasawara | Own | 15.4 | 143,041,238 |
| Ac95 | Ogasawara | Ogasawara | Own | 15.4 | 145,176,172 |
| Ac16 | Shikine | Mainland | Own | 16.4 | 151,977,572 |
| Ac17 | Shikine | Mainland | Own | 21.5 | 208,595,324 |
| Ac18 | Shikine | Mainland | Own | 18.0 | 170,409,890 |
| Ac19 | Shikine | Mainland | Own | 16.7 | 153,926,350 |
| Ac20 | Shikine | Mainland | Own | 17.4 | 163,354,786 |
| Ac21 | Shikine | Mainland | Own | 14.8 | 134,347,576 |
| Ac100 | Ishigaki | Ryukyu | Own | 17.8 | 165,763,146 |
| Ac102 | Ishigaki | Ryukyu | Own | 14.9 | 134,553,216 |
| Ac108 | Ishigaki | Ryukyu | Own | 16.3 | 151,398,928 |
| Ac110 | Ishigaki | Ryukyu | Own | 17.9 | 169,526,616 |
| Ac113 | Ishigaki | Ryukyu | Own | 16.1 | 151,566,068 |
| Ac115 | Ishigaki | Ryukyu | Own | 16.2 | 150,256,380 |
| Ac117 | Ishigaki | Ryukyu | Own | 17.6 | 162,985,410 |
| Ac126 | Ishigaki | Ryukyu | Own | 15.7 | 144,902,520 |
| Ac28 | Amami | Ryukyu | Own | 14.7 | 133,161,104 |
| Ac32 | Amami | Ryukyu | Own | 14.3 | 130,255,242 |
| Ac35 | Amami | Ryukyu | Own | 15.5 | 143,195,534 |
| Ac39 | Amami | Ryukyu | Own | 13.3 | 120,726,236 |
| Ac41 | Amami | Ryukyu | Own | 16.5 | 151,984,848 |
| Ac42 | Amami | Ryukyu | Own | 15.8 | 148,013,938 |
| Ac136 | Aka | Ryukyu | Own | 15.0 | 136,656,796 |
| Ac139 | Aka | Ryukyu | Own | 16.7 | 154,557,664 |
| Ac141 | Aka | Ryukyu | Own | 13.5 | 121,488,674 |
| Ac149 | Aka | Ryukyu | Own | 17.2 | 160,376,374 |
| Ac151 | Aka | Ryukyu | Own | 15.3 | 141,062,324 |
| Ac212 | Kagoshima | Mainland | Own | 17.4 | 161,010,430 |

| Sample name | Location | Population | Collection | Sequencing depth (mean) | Reads |
|---|---|---|---|---|---|
| Ac223 | Kagoshima | Mainland | Own | 17.5 | 163,857,650 |
| Ac235 | Kagoshima | Mainland | Own | 16.3 | 149,070,624 |
| Ac239 | Kagoshima | Mainland | Own | 17.0 | 156,570,028 |
| Ac244 | Kagoshima | Mainland | Own | 15.0 | 138,140,952 |
| Ac217 | Kagoshima | Mainland | Own | 21.7 | 150,424,554 |
| Ac221 | Kagoshima | Mainland | Own | 20.2 | 137,344,930 |
| Ac245 | Kagoshima | Mainland | Own | 24.4 | 167,324,940 |
| Ac266 | Yakushima | Mainland | Own | 16.3 | 151,426,358 |
| Ac276 | Yakushima | Mainland | Own | 15.4 | 144,332,374 |
| Ac277 | Yakushima | Mainland | Own | 17.4 | 164,038,776 |
| Ac288 | Yakushima | Mainland | Own | 15.8 | 146,484,300 |
| Ac289 | Yakushima | Mainland | Own | 21.3 | 205,309,868 |
| Ac263 | Yakushima | Mainland | Own | 30.7 | 213,721,582 |
| Ac264 | Yakushima | Mainland | Own | 26.0 | 181,366,660 |
| Ac3 | Okinawa | Ryukyu | Own | 17.3 | 160,675,234 |
| Ac169 | Okinawa | Ryukyu | Own | 18.4 | 175,948,500 |
| Ac183 | Okinawa | Ryukyu | Own | 19.8 | 186,513,930 |
| Ac197 | Okinawa | Ryukyu | Own | 15.7 | 144,751,218 |
| Ac297 | Okinawa | Ryukyu | Own | 18.5 | 178,771,772 |
| Ac179 | Okinawa | Ryukyu | Own | 24.9 | 173,446,608 |
| AcG2 | Guam | Guam | Own | 26.1 | 180,936,802 |
| AcG3 | Guam | Guam | Own | 24.0 | 163,697,788 |
| AcG4 | Guam | Guam | Own | 28.0 | 189,398,990 |
| SRR29150595 | Tulamben | Tulamben | NCBI † | 10.8 | 93,945,696 |
| SRR29150596 | Tulamben | Tulamben | NCBI † | 8.7 | 74,468,078 |
| SRR29150597 | Tulamben | Tulamben | NCBI † | 9.8 | 84,912,226 |
| SRR29150598 | Taiwan | Taiwan | NCBI † | 15.0 | 116,954,658 |
| SRR29150599 | Taiwan | Taiwan | NCBI † | 14.8 | 115,999,578 |
| SRR29150600 | Taiwan | Taiwan | NCBI † | 14.7 | 115,079,096 |
| SRR29150601 | Taiwan | Taiwan | NCBI † | 13.7 | 115,978,408 |
| SRR29150602 | Taiwan | Taiwan | NCBI † | 13.1 | 108,752,240 |
| SRR29150603 | Taiwan | Taiwan | NCBI † | 13.4 | 116,335,424 |
| SRR29150604 | Taiwan | Taiwan | NCBI † | 15.4 | 116,417,330 |
| SRR29150605 | Tulamben | Tulamben | NCBI † | 8.9 | 77,060,416 |
| SRR29150606 | Taiwan | Taiwan | NCBI † | 15.5 | 120,832,582 |
| SRR29150607 | Taiwan | Taiwan | NCBI † | 13.5 | 116,745,970 |
| SRR29150608 | Solomon | Solomon | NCBI † | 15.9 | 112,483,782 |
| SRR29150609 | Solomon | Solomon | NCBI † | 14.7 | 116,164,858 |
| SRR29150610 | Solomon | Solomon | NCBI † | 7.9 | 88,327,024 |
| SRR29150611 | Solomon | Solomon | NCBI † | 6.9 | 74,168,504 |
| SRR29150612 | Solomon | Solomon | NCBI † | 7.3 | 78,556,214 |
| SRR29150613 | Solomon | Solomon | NCBI † | 5.6 | 57,430,772 |
| SRR29150614 | Solomon | Solomon | NCBI † | 7.2 | 77,980,576 |
| SRR29150615 | Solomon | Solomon | NCBI † | 8.8 | 97,731,144 |

| Sample name | Location | Population | Collection | Sequencing depth (mean) | Reads |
|---|---|---|---|---|---|
| SRR29150616 | Tulamben | Tulamben | NCBI † | 9.0 | 79,046,136 |
| SRR29150617 | Solomon | Solomon | NCBI † | 6.9 | 74,590,936 |
| SRR29150618 | Philippines | Philippines | NCBI † | 10.9 | 98,104,966 |
| SRR29150619 | Philippines | Philippines | NCBI † | 14.0 | 116,245,758 |
| SRR29150620 | Philippines | Philippines | NCBI † | 14.2 | 114,409,430 |
| SRR29150621 | Philippines | Philippines | NCBI † | 15.4 | 118,376,834 |
| SRR29150622 | Philippines | Philippines | NCBI † | 15.6 | 118,465,290 |
| SRR29150623 | Philippines | Philippines | NCBI † | 14.4 | 116,781,168 |
| SRR29150624 | Philippines | Philippines | NCBI † | 14.5 | 116,637,138 |
| SRR29150625 | Philippines | Philippines | NCBI † | 14.1 | 117,565,750 |
| SRR29150626 | Philippines | Philippines | NCBI † | 14.7 | 116,210,520 |
| SRR29150627 | Manado | Manado | NCBI † | 11.9 | 112,240,892 |
| SRR29150628 | Philippines | Philippines | NCBI † | 14.9 | 119,043,898 |
| SRR29150629 | PNG | PNG | NCBI † | 14.3 | 117,749,588 |
| SRR29150630 | PNG | PNG | NCBI † | 14.1 | 115,607,374 |
| SRR29150631 | PNG | PNG | NCBI † | 12.9 | 116,181,130 |
| SRR29150632 | PNG | PNG | NCBI † | 13.9 | 113,941,722 |
| SRR29150633 | PNG | PNG | NCBI † | 14.7 | 116,262,726 |
| SRR29150634 | PNG | PNG | NCBI † | 13.3 | 111,851,402 |
| SRR29150635 | PNG | PNG | NCBI † | 12.7 | 111,108,952 |
| SRR29150636 | PNG | PNG | NCBI † | 12.4 | 112,327,741 |
| SRR29150637 | PNG | PNG | NCBI † | 13.3 | 113,351,863 |
| SRR29150638 | Manado | Manado | NCBI † | 11.3 | 105,311,982 |
| SRR29150639 | PNG | PNG | NCBI † | 13.2 | 113,470,477 |
| SRR29150640 | PNG | PNG | NCBI † | 13.5 | 113,431,547 |
| SRR29150641 | NewCal | NewCal | NCBI † | 15.4 | 119,252,088 |
| SRR29150642 | NewCal | NewCal | NCBI † | 13.7 | 116,733,916 |
| SRR29150643 | NewCal | NewCal | NCBI † | 9.1 | 80,712,980 |
| SRR29150644 | NewCal | NewCal | NCBI † | 9.7 | 86,448,912 |
| SRR29150645 | NewCal | NewCal | NCBI † | 10.8 | 95,932,054 |
| SRR29150646 | NewCal | NewCal | NCBI † | 11.3 | 101,024,234 |
| SRR29150647 | NewCal | NewCal | NCBI † | 9.7 | 86,803,052 |
| SRR29150648 | NewCal | NewCal | NCBI † | 13.5 | 114,203,692 |
| SRR29150649 | Manado | Manado | NCBI † | 9.7 | 90,384,846 |
| SRR29150650 | Maldives | Maldives | NCBI † | 9.4 | 83,135,924 |
| SRR29150651 | Maldives | Maldives | NCBI † | 9.1 | 80,089,302 |
| SRR29150652 | Maldives | Maldives | NCBI † | 11.9 | 106,539,814 |
| SRR29150653 | Maldives | Maldives | NCBI † | 10.7 | 93,940,056 |
| SRR29150654 | Maldives | Maldives | NCBI † | 9.6 | 84,866,426 |
| SRR29150655 | Maldives | Maldives | NCBI † | 8.3 | 78,791,708 |
| SRR29150656 | Maldives | Maldives | NCBI † | 13.3 | 127,495,358 |
| SRR29150657 | Maldives | Maldives | NCBI † | 11.0 | 107,566,094 |
| SRR29150658 | Maldives | Maldives | NCBI † | 12.0 | 113,849,868 |
| SRR29150659 | Maldives | Maldives | NCBI † | 11.3 | 109,091,406 |

| Sample name | Location | Population | Collection | Sequencing depth (mean) | Reads |
|---|---|---|---|---|---|
| SRR29150660 | Manado | Manado | NCBI † | 10.6 | 100,537,870 |
| SRR29150661 | Manado | Manado | NCBI † | 11.2 | 104,562,370 |
| SRR29522765 | Marshall Island | *A. tricinctus* | NCBI † | 8.6 | 99,952,268 |
| SRR31951262 | na | *A. chrysopterus* | NCBI † | 5.9 | 44,488,128 |

† NCBI accession number is same as sample name.

[1] From: García Jiménez, A., Talbi, M., Fitzgerald, L. M., Heim, A., Marcionetti, A., Schmid, S., Bertrand, J., Shaughnessy, A., Santiago, C., & Rangseethampanya, P. (2025). Habitat Specialisation Impacts Clownfish Demographic Resilience to Pleistocene Sea-Level Fluctuations. *Molecular Ecology*, e70134. https://doi.org/10.1111/mec.70134

**Table S2:** Individual pairs with a kinship coefficient larger than 0.088, indicating 2nd degree relatives. ID1 and ID2: the individual sample names; N_SNP: Number of SNPs used in the pairwise analysis.

| Population | ID1 | ID2 | N_SNP | Kinship coefficient |
|---|---|---|---|---|
| Thailand | CLK_113 | CLK_120 | 35153023 | 0.1388 |
| Thailand | CLK_150 | CLK_76 | 35126390 | 0.1276 |
| Guam | AcG3 | AcG4 | 34945136 | 0.1249 |
| Thailand | CLK_123 | CLK_133 | 35156324 | 0.1228 |
| Guam | AcG2 | AcG3 | 34951417 | 0.1187 |
| Thailand | CLK_133 | CLK_150 | 35147658 | 0.1185 |
| Guam | AcG2 | AcG4 | 34950309 | 0.116 |
| Thailand | CLK_123 | CLK_130 | 35146182 | 0.1157 |
| Thailand | CLK_130 | CLK_133 | 35158510 | 0.1152 |
| Thailand | CLK_123 | CLK_150 | 35136398 | 0.1147 |
| Thailand | CLK_113 | CLK_130 | 35155372 | 0.1133 |
| PNG | SRR29150629 | SRR29150634 | 35091953 | 0.113 |
| PNG | SRR29150632 | SRR29150639 | 35082578 | 0.113 |
| Thailand | CLK_113 | CLK_123 | 35151914 | 0.1129 |
| PNG | SRR29150632 | SRR29150634 | 35084779 | 0.1127 |
| Thailand | CLK_113 | CLK_133 | 35165664 | 0.1127 |
| PNG | SRR29150629 | SRR29150632 | 35095113 | 0.1116 |
| PNG | SRR29150635 | SRR29150639 | 35073297 | 0.1114 |
| PNG | SRR29150632 | SRR29150633 | 35094701 | 0.1111 |
| Thailand | CLK_123 | CLK_91 | 35159682 | 0.1111 |
| Thailand | CLK_133 | CLK_91 | 35172817 | 0.1107 |
| Thailand | CLK_150 | CLK_91 | 35149975 | 0.1099 |
| PNG | SRR29150634 | SRR29150639 | 35080538 | 0.1092 |
| Thailand | CLK_113 | CLK_150 | 35143262 | 0.109 |
| PNG | SRR29150632 | SRR29150635 | 35076933 | 0.1081 |
| Thailand | CLK_130 | CLK_150 | 35136923 | 0.1073 |
| PNG | SRR29150633 | SRR29150634 | 35089897 | 0.1072 |
| PNG | SRR29150629 | SRR29150635 | 35082296 | 0.1061 |
| PNG | SRR29150634 | SRR29150635 | 35073161 | 0.1058 |
| PNG | SRR29150633 | SRR29150635 | 35083034 | 0.1041 |
| PNG | SRR29150633 | SRR29150639 | 35089931 | 0.1013 |
| PNG | SRR29150629 | SRR29150639 | 35089389 | 0.1007 |
| PNG | SRR29150629 | SRR29150633 | 35099978 | 0.1004 |
| Thailand | CLK_113 | CLK_91 | 35169815 | 0.099 |
| THA | CLK_130 | CLK_91 | 35161378 | 0.0956 |
| SOL | SRR29150611 | SRR29150612 | 34542234 | 0.0892 |

**Table S3:** Summary of Runs of Homozygosity (RoH) for each population. #Segments: Average number of RoH per population ± standard deviation; KBSegment: Average length of the RoH in kilobases ± standard deviation.

| | #Segments | KBSegment |
|---|---|---|
| ***A. tricinctus*** | 108.0 ± 6.2 | 407.4 ± 13.6 |
| **Bali** | 19.0 ± 3.7 | 398.0 ± 20.7 |
| **Guam** | 423.3 ± 20.3 | 440.6 ± 6.7 |
| **Japan** | 12.9 ± 6.0 | 411.0 ± 45.9 |
| **Maldives** | 64.7 ± 7.9 | 399.9 ± 11.8 |
| **New Caledonia** | 176.3 ± 15.6 | 408.3 ± 6.0 |
| **Ogasawara** | 208.2 ± 11.1 | 417.3 ± 8.9 |
| **Philippines** | 14.0 ± 3.7 | 423.0 ± 23.3 |
| **Papua New Guinea** | 239.5 ± 11.6 | 405.2 ± 7.1 |
| **Ryukyus** | 13.9 ± 5.3 | 413.2 ± 35.3 |
| **Solomon Islands** | 246.9 ± 69.3 | 411.6 ± 21.3 |
| **Sulawesi** | 15.8 ± 1.3 | 423.8 ± 19.0 |
| **Taiwan** | 39.3 ± 16.1 | 451.6 ± 34.8 |
| **Thailand** | 62.4 ± 16.0 | 427.2 ± 29.0 |
